# Sex-specific prefrontal cortex dysfunction underlying opioid-induced cognitive impairment

**DOI:** 10.1101/666099

**Authors:** Eden M. Anderson, Annabel Engelhardt, Skyler Demis, Elissa Porath, Matthew C. Hearing

## Abstract

Women transition to addiction faster and experience greater difficulties remaining abstinent; however, what drives this is unknown. Although poorly understood, loss of cognitive control following chronic drug use has been linked to decreased activation of frontal cortical regions. We show that self-administration of the opioid, remifentanil, causes a long-lasting decrease in *ex vivo* excitability but augments firing capacity of pyramidal neurons in the prelimbic cortex. This phenomenon occurs faster in females, manifests from sex-specific changes in excitatory and inhibitory synaptic regulation and aligns with impairments in cognitive flexibility. Further, chemogenetic induction of a hypoactive pyramidal neuron state in drug-naïve mice produces deficits, while compensating for this hypoactive state protects against cognitive inflexibility resulting from opioid self-administration. These data define cellular and synaptic mechanisms by which opioids impair prefrontal function and cognitive control and indicate that interventions aimed at treating opioid addiction must be tailored based on biological sex.

## Introduction

Even when taken as prescribed, use of opioid-based drugs carry a risk of misuse and dependence that can lead to out of control drug use. Although opioid use disorders are prevalent among men and women, women report more lifetime use of prescription opioids^1^, greater opioid craving, experience greater difficulties remaining abstinent^2,3^, transition to compulsive drug use faster, and are at greater risk for overdose related deaths4. While environmental and psychological factors may contribute to these discrepancies, rodent models of addiction exhibit similar trends with females showing more rapid acquisition of heroin-taking behavior and greater overall drug intake^5,6^, indicating that the neural mechanisms underlying the transition to opioid addiction is distinct across sex.

Deficits in cognitive control, whether intrinsic or arising from early or prolonged drug use, increase the risk and severity of addiction^7,8^. Although the range of cognitive problems in addiction is diverse, the most consistently documented deficits include cognitive inflexibility and reduced inhibitory control^8–10^. Clinical data indicates that deficits in the ability to adapt one’s behavior in response to changing environmental contingencies (i.e. cognitive flexibility) can increase the risk and severity of addiction by strengthening drug seeking behaviors, increasing relapse vulnerability, and impair an individual’s ability to resist habitual drug use^7,11–15^. Thus, identifying adaptations responsible for impaired decision-making that precede and/or parallel out of control drug use provides an opportunity to increase efficacy of treatments aimed at mitigating relapse and reducing susceptibility for developing opioid use disorders.

The prelimbic region of the medial PFC (mPFC) in rodents is known to govern numerous cognitive functions, including encoding of flexible decision-making, inhibitory control, and opioid-seeking^7,16–19^. Impaired cognition in numerous pathological states has been linked to reduced or increased/irregular spike firing activity in the prelimbic cortex^20–23^. Clinical imaging studies have identified a dichotomous dysfunction of an area homologous to the prelimbic in individuals addicted to heroin, showing reduced basal levels of metabolic activity as well as craving related increases in neural activity in response to drug cues^24–28^. Data from rodent addiction models has separately shown a similar phenomenon whereby prelimbic neurons exhibit a progressive enhancement in their response to cocaine and cocaine cues, while reduced excitability of prelimbic layer 5/6 (L5/6) pyramidal neurons following extended cocaine exposure promotes compulsive drug-seeking^7,16,18,29–34^. Compared to drugs of abuse such as cocaine, far less is known regarding the impact of opioid exposure on PFC cellular physiology and synaptic regulation, with even less known about how biological sex confers vulnerability to these adaptations. Here, we outline time- and sex-specific effects of opioid self-administration on intrinsic excitability and synaptic regulation of pyramidal neurons in the prelimbic and infralimbic region of the mPFC, and the resultant impact on cognitive flexibility in both males and females.

## Results

### Effects of short- and long-term remifentanil self-administration on layer 5/6 prelimbic pyramidal neurons

To determine how contingent administration of a clinically relevant opioid impacts PFC function, male and female mice originally underwent 14 days of saline or remifentanil (Ultiva®) self-administration (short-term) followed by 14-21 days of forced abstinence (Figure 1a). During initial studies, mice were lever trained using a liquid Ensure® reward on a fixed ratio (FR1) schedule (not paired), followed by 14 days of intravenous (i.v.) saline or remifentanil self-administration (3 hours/day). Subsequent studies utilized a training approach whereby i.v. infusions were initially paired with delivery of Ensure® on an FR1-FR2-FR3 schedule, followed by i.v. administration without Ensure® during 2 hour daily sessions. This was done to expedite acquisition and reduce variability in total days of drug exposure (i.e., reduce days below criterion), reduce the rate of failed acquisition, and increase daily throughput. When Ensure® was not paired during training, remifentanil mice had significantly more active lever presses compared to saline mice (Supplemental 1a), whereas pairing with Ensure® prompted a sustained increase in active lever responding in saline mice (Supplemental 1b), negating an overall difference across drug treatment (Supplemental 1c). Active lever responding and cumulative infusions remained similar in paired and not paired remifentanil mice (Supplemental 1d-e). Moreover, paired male remifentanil mice exhibited higher break points for remifentanil compared to paired saline mice for a saline infusion (Supplemental 1f), indicating that the selected dose is reinforcing. Comparison of 2 and 3 hour daily sessions showed that total infusions earned in remifentanil males and females did not differ nor did the overall number of active or inactive presses (Supplemental 2); thus data from these initial cohorts were combined.

**Figure 1.**
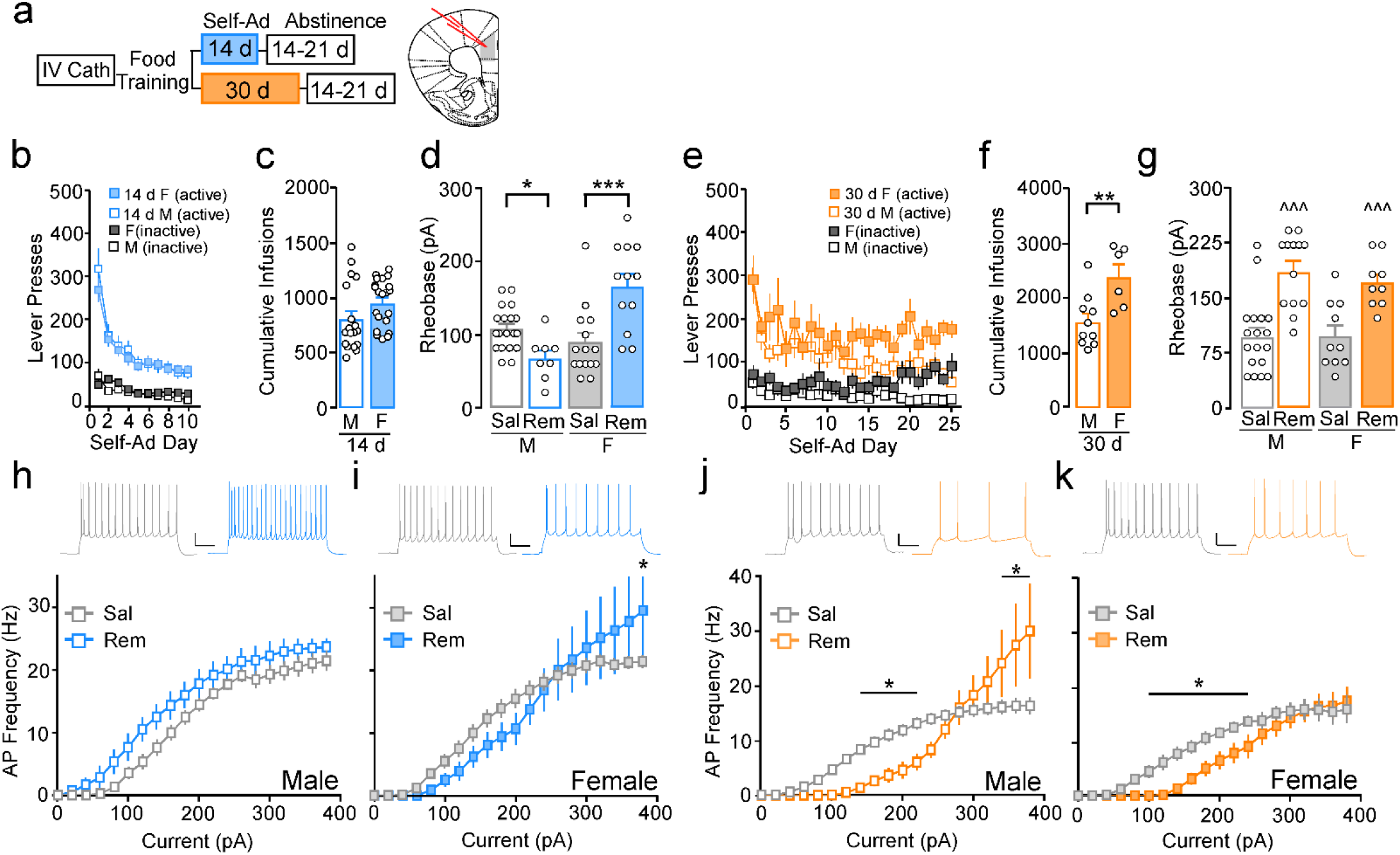
Bidirectional effect of remifentanil self-administration on prelimbic L5/6 pyramidal neurons in males and females. **a,** Self-administration, abstinence timeline, and location of whole-cell recordings in the prelimbic cortex. **b,** For mice that received 14 days of remifentanil self-administration (short-term), no sex by day interaction (F_(9,333)_= 0.68, p=0.73) or main effect of sex (F_(1,37)_= 0.06, p=0.81) was observed for active presses. For inactive lever presses, a significant day by sex interaction (F_(9, 333)_= 1.91, p=0.05) was observed with post-hoc comparisons showing males having lower presses on day 2 (p=0.005). **c,** Cumulative remifentanil infusions did not differ in males (M) versus females (F) (t_(37)_= −1.69, p=0.10). **d,** Sex by treatment interaction (F_(1,50)_= 22.87, p<0.001) comparing mean action potential (AP) threshold (rheobase) in prelimbic L5/6 pyramidal neurons following 14-21 d abstinence from short-term self-administration. Rheobase was similar in females compared to males under saline conditions (SAL) (N/n= F: 9/15, M:13/19, p=0.22). Remifentanil (REM) increased rheobase in females (N/n=4/12, p<0.001) but reduced it in males (N/n=6/8, p=0.037) compared to respective controls. **e,** For long-term self-administration groups, no sex by day interaction (F_(24,336)_= 1.07, p=0.37) or main effect of sex (F_(1,14)_= 3.67, p=0.08) on active presses was observed. There was a sex by day interaction on inactive presses (F_(24, 336)_= 1.78, p=0.02), with males having significantly lower inactive presses on days 3, 9, 13 (p<0.05), 19-20 (p=0.001), 21 (p<0.05), 22 (p<0.01), 23 (p<0.001) and 24 (p<0.01). **f,** Cumulative remifentanil infusions were elevated in females (F) compared to males (M) (t_(14)_=−3.12, p=0.008). **g,** Comparison of mean rheobase following 14-21 d abstinence from long-term self-administration showed an interaction of sex by treatment (F_(1,46)_= 34.13, p<0.001), with remifentanil males (N/n=6/13) and females (N/n=4/9) exhibiting higher firing thresholds compared to respective saline controls (Male: N/n=9/18, Female: N/n=5/10). **h-i,** Current-spike analysis in short-term self-administering mice showed a trend toward increased firing frequency in remifentanil (N/n=6/13) versus saline (N/n=9/18) males (**h,** *treatment*: F_(1,19)_=3.78, p=0.07) and no significant interaction (F_(19,361)_= 0.62, p=0.90), but an interaction of treatment by current in females (**i**, F_(19,418)_=1.61, p=0.05; N/n= Sal:9/14, Rem:4/10). **j,** Following long-term self-administration, current-spike analysis showed firing frequency was reduced at lower currents but increased at more depolarized potentials in remifentanil (N/n=6/13) versus saline (N/n=9/18) males (*interaction*: F_(19,551)_=5.43, p<0.001). **k,** Remifentanil females (N/n=4/9) showed reduced firing frequency at lower currents but similar firing frequency at more depolarized potentials versus saline females (N/n=5/10; interaction: F_(19,323)_= 2.22, p=0.003). Representative scale bar: 20pA/200msec. All data are presented as the mean ± SEM. *p<0.05, **p<0.01, ***p<0.001.

Functional integrity of mPFC information processing is dependent on extrasynaptic factors that dictate pyramidal neuron firing threshold and responsivity to synaptic input^35–38^. Thus, we first used *ex vivo* electrophysiology to examine these properties in L5/6 prelimbic pyramidal neurons following short-term remifentanil self-administration. During short-term self-adminsitration, active lever responding and cumulative infusions earned did not vary across males and females (Figure 1b-c). Following 14-21 days of abstinence, intrinsic excitability of L5/6 prelimbic pyramidal neurons was assessed by measuring the threshold to evoke an initial action potential (rheobase) in response to depolarizing current steps (1 s, 0-380 pA, 20 pA steps). In males, remifentanil reduced mean rheobase compared to saline males; however, in females, firing threshold was increased (Figure 1d). As cell excitability reflects a combination of firing threshold and spike activity states, and rheobase is more defined as a measure of basal excitation, we will refer to reductions and increases in threshold as hyperactive and hypoactive basal states, respectively. These adaptations were not influenced by extinction from initial Ensure® training, as additional cohorts undergoing Ensure® self-administration for ~14 days showed similar rheobase compared to those undergoing subsequent saline administration and remained different from rheobase in remifentanil mice (Supplemental 3).

To determine whether this hypoactive basal state is unique to females or merely arises on a more rapid time scale compared to males, rheobase was measured from mice who administered saline or remifentanil for 30 days (long-term). Similar to short-term self-administration, active lever pressing did not vary (Figure 1e); however, females showed greater intake compared to their male counterparts (Figure 1f). Following a similar abstinence period, assessment of rheobase showed that female, and now male, remifentanil mice exhibited an increase in mean threshold to fire an action potential compared to controls (Figure 1g). These data demonstrate that similar to past findings with cocaine in males, opioid self-administration promotes a hypoactive basal state in prelimbic pyramidal neurons, however, this dysfunction arises more rapidly in females.

Dysfunction of frontal cortex and impaired cognitive control has been linked to irregularities in PFC pyramidal neuron firing patterns across numerous disorders^20–23^. Thus, we next examined how opioid self-administration impacted patterned spike firing. Assessment of current-spike relationships of pyramidal neurons in males following short-term remifentanil self-administration exhibited a nonsignificant leftward shift and increase in firing frequency compared to saline controls (Figure 1h). Alternatively, pyramidal neurons in females aligned with reduced firing at lower currents, but increased firing at more depolarized potentials (Figure 1i). As action potential firing plateaus or decreases at these higher potentials in control conditions, it suggests hypoactive basal states are paralleled by an increased capacity to maintain firing once activated. Following long-term self-administration, pyramidal neurons in males showed a similar divergent shift in firing akin to that observed in females following short-term self-administration (Figure 1j). This shift was no longer present in females, with firing frequency reduced overall versus saline controls (Figure 1k). Notably, no significant effects on excitability or firing were observed in infralimbic cortex pyramidal neurons following short- or long-term self-administration (Supplemental 4).

Previous studies with cocaine in males indicate that the prelimbic cortex undergoes time- dependent alterations in activity and plasticity-related gene expression^31,39–46^. To further characterize the temporal nature of opioid-induced adaptations, we first asked whether hyperexcitable states observed in males after short-term self-administration occurs in females, but on a shorter time scale. Following 5 days of self-administration and 14-21 day abstinence, mean rheobase was reduced in remifentanil male mice but increased in females (Figure 2a), despite males and females having similar responding and intake throughout self-administration (Supplemental 5). Similar to short-term exposure, changes in rheobase aligned with a leftward and rightward shift in action potential frequency in males and females, respectively (Figure 2b-c).

**Figure 2.**
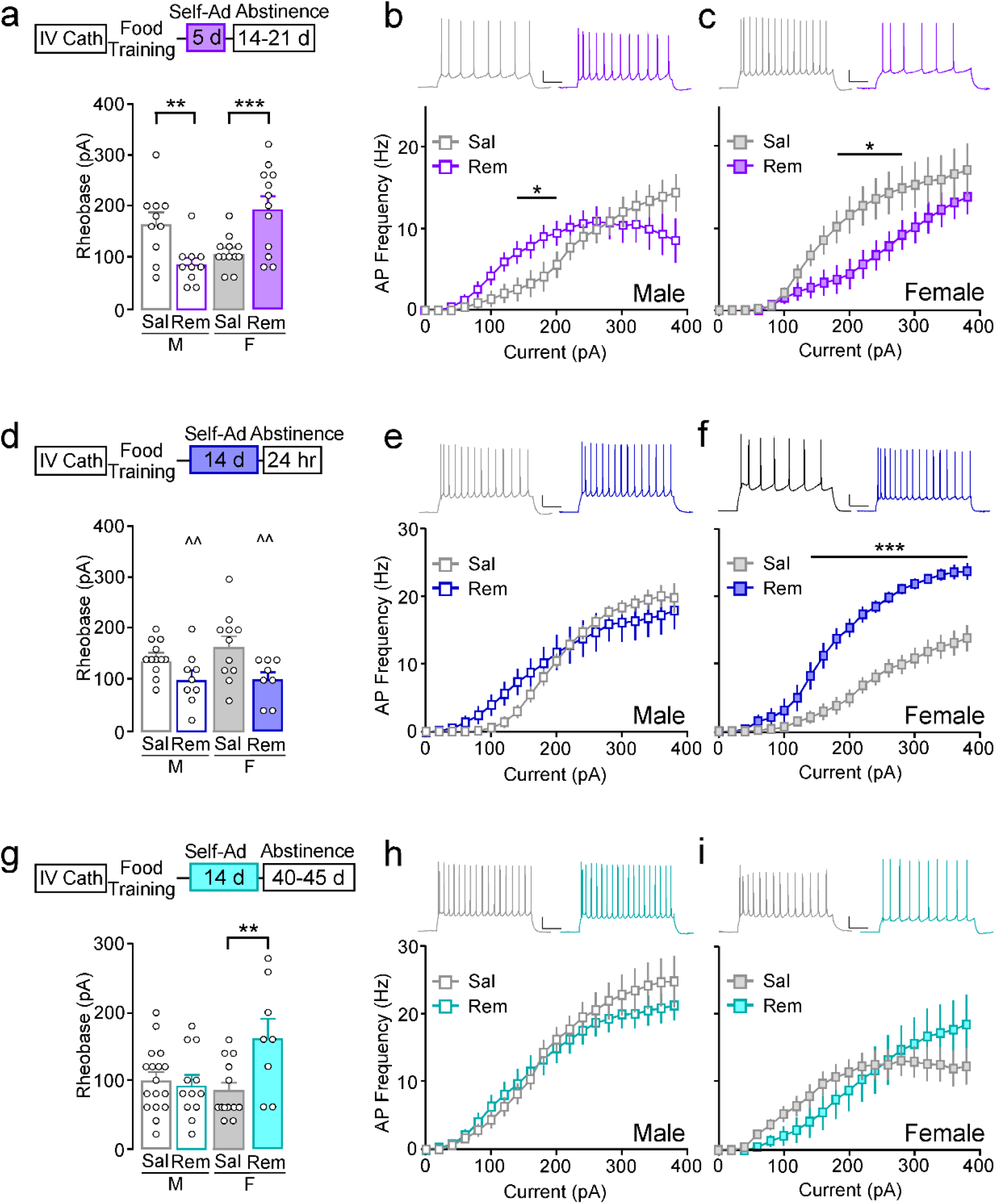
Time course of remifentanil self-administration effects on prelimbic L5/6 pyramidal neurons action potential threshold and firing. **a,** Self-administration and abstinence timeline. Following 5 days of self-administration and 14-21 days abstinence, an interaction of sex by treatment was observed for rheobase (F_(1, 40)_= 21.01, p<0.001). Rheobase was reduced in remifentanil males (N/n=4/10) compared to saline males (N/n=4/10; p=0.005) and increased in remifentanil females (N/n=4/12) compared to saline females (N/n=4/12; p=0.001). **b-c,** Current-spike analysis show a treatment by current interaction in both males (F_(19,342)_= 2.77, p<0.001) and females (F_(19,418)_= 2.45, p<0.001). Post-hoc comparisons show that remifentanil males (N/n=4/10) have greater firing at 140-200pA compared to saline males (N/n=4/10; p<0.05) whereas remifentanil females (N/n=4/12) have reduced firing at 180-280pA compared to saline females (N/n=4/12; p<0.05). **d.** Following 14 days of remifentanil with 24-72 hr abstinence, there was main effect of treatment with prelimbic L5/6 pyramidal exhibited lower rheobase in remifentanil males (N/n=5/9) and females (N/n=3/8) versus respective saline controls (M: N/n=4/12, F: N/n=4/11; F_(1,36)_= 10.77, p=0.002). **e-f,** Current-spike analysis show a treatment by current interaction in males (F_(19, 361)_= 2.62, p<0.001) and females (F_(19, 323)_= 8.95, p<0.001). Post-hoc comparisons showed no difference in remifentanil (N/n=5/9) versus saline males (N/n=4/12), whereas spike frequency was elevated in remifentanil females (N/n=4/12) at 140-380pA versus saline (N/n=4/12; p<0.001). **g,** Following 45 day abstinence from 14 days of self-administration, a significant sex by treatment interaction was observed for rheobase (F_(1,46)_= 7.38, p=0.009). Rheobase in males did not differ (Sal: N/n=10/17, Rem: N/n=7/12, p=0.70), while rheobase remained increased in remifentanil females (N/n=4/8) versus saline (N/n=7/13, p=0.002). **h-i,** Current-spike analysis showed no difference in firing frequency in remifentanil males (N/n=7/12) compared to saline (N/n=9/15; *interaction*: F_(19,475)_= 0.78, p=0.74). A significant treatment by current interaction was found in females (F_(19,342)_= 2.70, p<0.001), with a trend towards increased firing at 360-380pA in remifentanil mice (N/n=4/8) compared to saline (N/n=5/12; p<0.10). Representative scale bar: 20pA/200msec. All data are presented as the mean ± SEM. *p<0.05, **p<0.01, ***p≤0.001, ^^p<0.01 main effect of treatment.

To determine the timeline in which these adaptations occur following drug exposure, we examined alterations in excitability 24-72 hours or 40-45 days following short-term self-administration. Following acute abstinence, remifentanil mice showed reduced rheobase compared to respective saline controls (Figure 2d). Current-spike relationships showed no difference in firing frequency in males, while remifentanil increased firing frequency in females (Figure 2e-f). After prolonged abstinence, hypoactive basal states and elevated firing capacity previously observed following 14-21 days of abstinence remained present in females; however, reductions in firing threshold and increased firing previously observed in males was no longer present (Figure 2g). Together, these data highlight the enduring nature of hypoactive basal states in females and suggest that pyramidal neurons in females and males may undergo a unique time-dependent shift in excitability during abstinence.

### Impact of remifentanil on prelimbic inhibitory and excitatory synaptic regulation

Activation and firing patterns of pyramidal neurons are heavily influenced by synaptic and perisynaptic excitatory and inhibitory signaling at the soma and dendrites^36,37^. We have previously shown that cocaine-induced hyperexcitablity of prelimbic pyramidal neurons is linked to reductions in GABA_B_R-dependent activation of G protein inwardly-rectifying K+ channels (GABA_B_R-GIRK)^41^. Thus, we examined if changes in excitability following short- or long-term opioid exposure also aligned with altered GABA_B_R-GIRK signaling (Figure 3a). Somatodendritic currents evoked by the GABA_B_R agonist, baclofen (I_Baclofen_; 200 μM), were similar in short- and long-term saline males and females and thus were combined into a single saline group. In males, I_Baclofen_ was reduced following short-term self-administration, but increased following long-term self-administration compared to saline controls (Figure 3b-c), thus paralleling a decrease and increase in rheobase, respectively. Conversely, I_Baclofen_ was not altered by short- or long-term self-administration in females. Similar to changes in rheobase and spike firing in males, alterations in I_Baclofen_ were no longer present in males at 45 days abstinence Supplemental 6).

**Figure 3.**
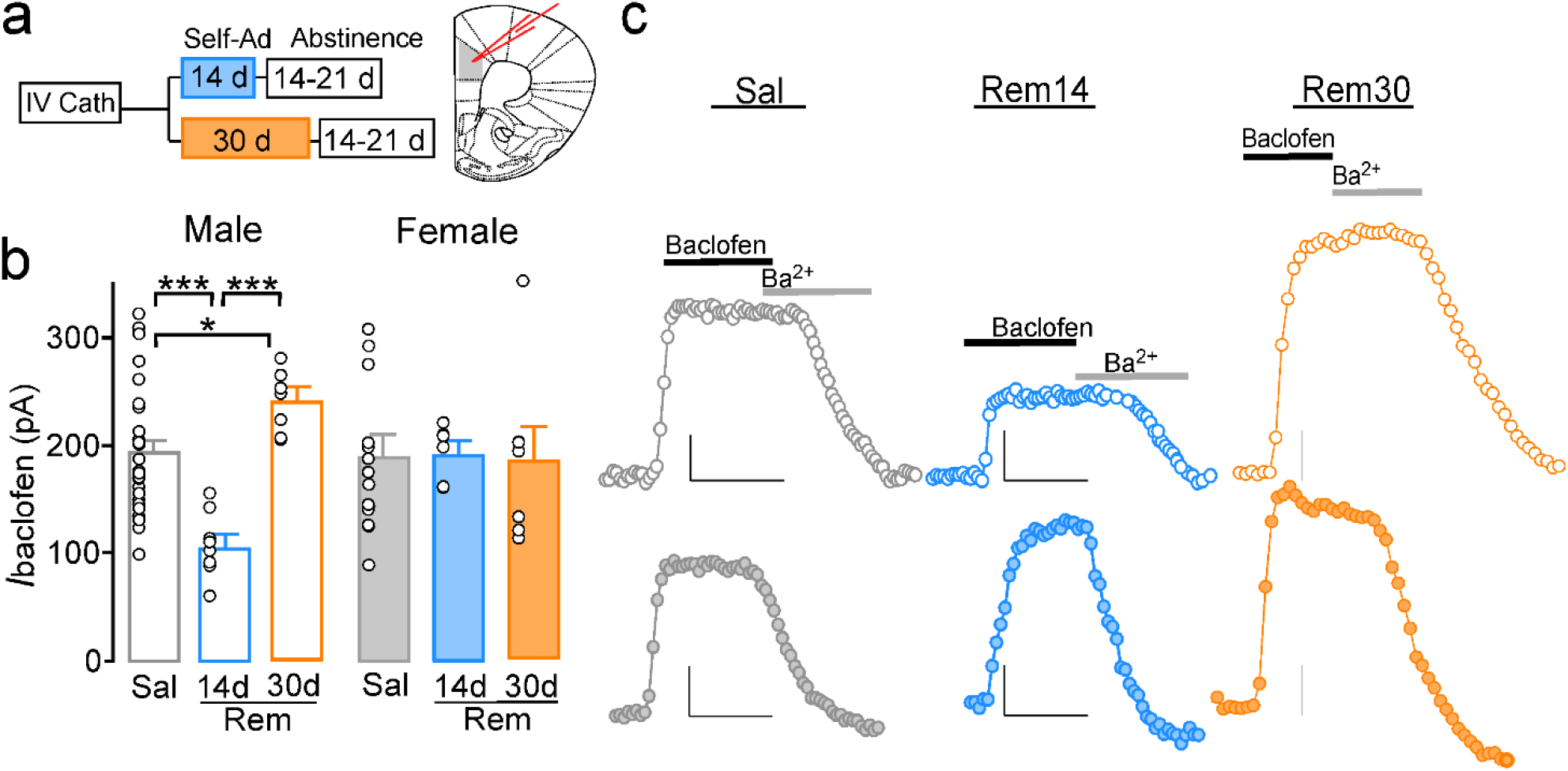
Remifentanil effects on prelimbic L5/6 pyramidal neurons GABA_B_-GIRK signaling. **a,** Self-administration and abstinence timepoints. **b,** Mean baclofen-evoked currents (*I*_baclof en_) from male (unfilled) and female (filled) prelimbic L5/6 pyramidal neurons 14-21 days after saline (gray), short-term remifentanil self-administration (blue) or long-term remifentanil self-administration (orange). *I*_baclof en_ was no different in saline short- or long-term self-administration males (14 d: N/n=11/13, 30 d: N/n=8/12; t_(22)_=0.442, p=0.663) or females (14 d: N/n=5/5, 30 d: N/n=7/9; t_(12)_=−0.842, p=0.416) and thus were combined. A significant sex by treatment interaction was observed (F_(2,60)_=5.79, p=0.005). *I*_baclof en_ was reduced in short-term remifentanil males (N/n=6/8) versus saline males (p<0.001) and long-term remifentanil males (N/n=6/7, p<0.001), whereas long-term remifentanil males showed an increase in *I*_baclof en_ versus saline males (p=0.037). *I*_baclof en_ did not differ when comparing saline females with short- (N/n=4/6, p=0.913) or long-term remifentanil self-administration (N/n=5/7, p=0.868). A difference was observed between short-term remifentanil males and females (p=0.006). **c,** Representative *I*_Baclof en_ traces in males (**top**) and females (**bottom**) across treatments. Representative scale bars: 50pA/200sec. *p<0.05, ***p<0.001.

Given the lack of change in GABA_B_R-GIRK signaling in females, we next examined modifications in ionotropic excitatory and inhibitory synaptic transmission by measuring AMPA receptor (AMPAR)-specific miniature excitatory postsynaptic currents (mEPSCs) and GABA_A_R-specific inhibitory postsynaptic currents (mIPSCs). Saline groups were again not different on any measure and were therefore combined (Supplemental 7). In females, mEPSC frequency was reduced in short- and long-term remifentanil mice compared to saline controls (Figure 4c). While remifentanil did not alter mEPSC frequency in males, baseline mEPSC frequency was higher in saline females compared to saline males. Only a main effect of treatment was identified for mEPSC amplitudes, with frequency elevated in long-term remifentanil mice compared to short-term remifentanil and saline groups (Figure 4d). Assessment of mIPSCs showed no differences in frequency across treatments in males (Figure 4g). mIPSC frequency was elevated in females following short-term remifentanil administration; however, these effects were no longer present following more prolonged exposure (Figure 4g). A main effect of sex was observed for mIPSC amplitude, with mean amplitudes greater in females compared to males, regardless of treatment (Figure 4h). These data indicate that alterations in pyramidal neuron excitation in males reflect divergent synaptic modifications following short- and long-term exposure, and that hypoactive basal states are driven via distinctly different mechanisms in males and females.

**Figure 4.**
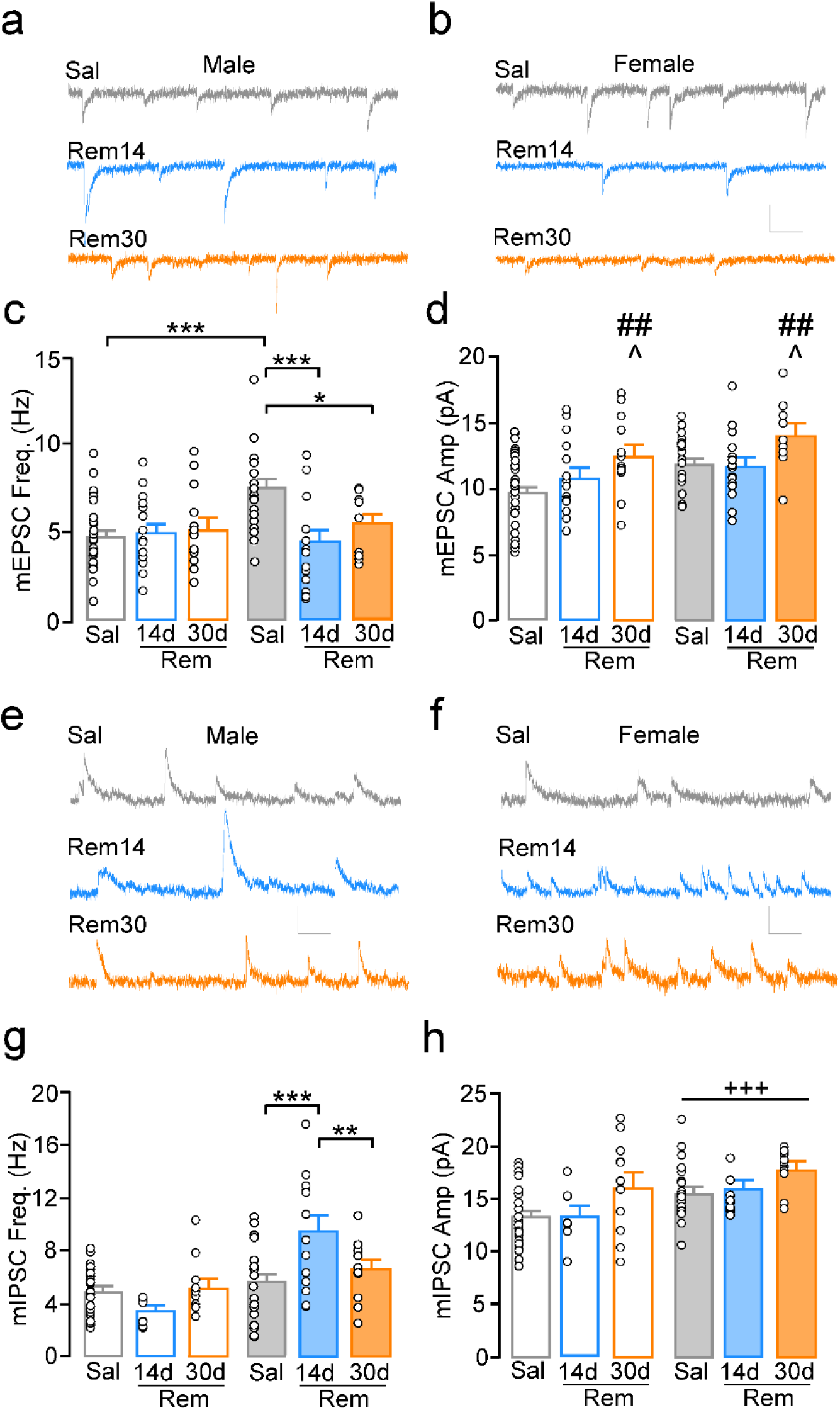
Remifentanil effects on prelimbic L5/6 pyramidal neuron excitatory and inhibitory transmission. **a-b,** Representative miniature excitatory postsynaptic current (mEPSC) current traces from prelimbic L5/6 pyramidal neurons in **(a)** males and **(b)** females following 14-21 days abstinence from self-administration. **c,** Comparison of mean mEPSC frequency identified an interaction of sex by treatment (F_(1,94)_=4.66, p=0.01). Saline females exhibited elevated mean frequenc y compared to 14 day (N/n=8/15, p=0.001) and 30 day (N/n=4/9, p=0.038) remifentanil females and compared to saline males (p<0.001). Mean mEPSC frequency did not differ when comparing saline males with 14 day (N/n=8/15, p=0.73) or 30 day (N/n=5/13, p=0.79) remifentanil males. **d,** Comparison of mEPSC amplitude showed a significant main effect of treatment, but not sex or a sex by treatment interaction (*treatment*: F_(2,94)_=4.77, p=0.01; sex: F_(1,94)_=3.63, p=0.06; interaction: F_(2,94)_=10.27, p=0.76). Effects within treatment showed increased mean mEPSC amplitude in long-term remifentanil mice compared to short-term remifentanil (p=0.019) and saline (p=0.006) mice independent of sex. **e-f,** Representative miniature inhibitory PSC (mIPSC) traces 14-21 days following short-term **(e)** and long-term **(b)** self-administration. **g,** Comparison of mean mIPSC frequency identified a sex by treatment interaction (F_(2,92)_=7.31, p=0.001). Mean frequency was increased in short-term remifentanil females (N/n=10/13) versus saline females (p<0.001) and long-term remifentanil females (N/n=4/11; p=0.009), with no differences between long-term remifentanil versus saline females (p=0.33). No differences were observed when comparing saline males to short- (N/n=5/8; p=0.19) or long-term (N/n=5/11; p=0.67) remifentanil males. **h,** A main effect of sex, but not treatment, or interaction was observed for mIPSC amplitude (*sex*: F_(1,87)_=11.83, p<0.001; *treatment*: F_(1,87)_=1.73, p=0.18; *interaction*: F_(1,87)_=0.22, p=0.81), with females showing higher amplitudes compared to males, regardless of treatment. Scale bars: 10pA/100msec. *p<0.05, **p<0.01, ***p<0.001; ^+++^p<0.001 main effect of sex, ^p<0.05 vs Rem14, ^##^p<0.01 vs Rem30.

### Time-dependent effects of remifentanil on affect and cognitive flexibility

Prelimbic dysfunction has been linked to impairments in flexible behavior^17,47^ and affect dysregulation that are known to increased risk for relapse and heightened drug use^7,16,28,48–50^. We first examined whether short-term opioid self-administration produced changes in performance in the elevated plus maze (EPM) and a forced swim test (FST) following short-term remifentanil self-administration, as this time point identified sex-differences in plasticity. Percent open arm time in the EPM was not altered in male or females following acute abstinence; however, females showed an increase in time immobile in the FST following more prolonged abstinence (Supplemental 8).

To examine effects on cognitive flexibility, we used an operant-based attentional set-shifting model following short-term self-administration. This task resembles the Wisconsin Card Sorting Task in the sense that the stimuli are easy to detect, but rules are implicit and learned while the task is performed (Figure 5a). Further, it has particular relevance to addiction, as chronic drug users exhibit attentional bias that make it more difficult to change behavior^9,10,51–54^. Remifentanil exposure did not alter days to reach lever training criteria (Figure 5b) or performance during the visual cue discrimination task, indicating that discriminative and associative learning processes remain intact in males and females (Figure 5c; Supplemental 9a). Alternatively, remifentanil females required significantly more trials and had more errors to reach criterion during an extradimensional shift test compared to saline females, whereas no differences were observed in males (Figure 5d; Supplemental 9b). Performance during a subsequent reversal learning test -- a task more reliant on the orbitofrontal region^55^ -- was also unaffected in both males and females (Figure 5e; Supplemental 9c).

**Figure 5.**
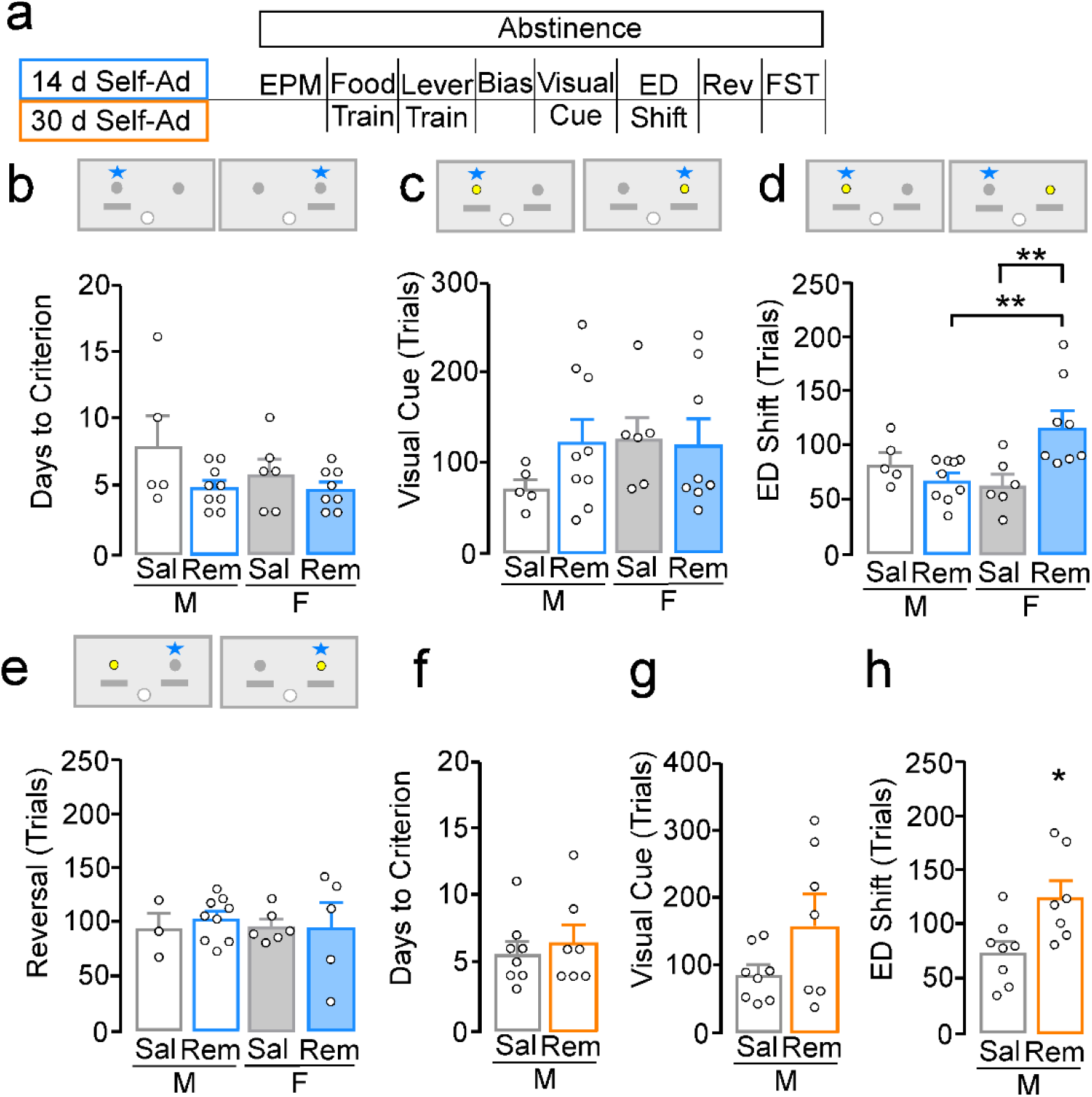
Remifentanil-induced deficits in cognitive flexibility. **a,** Timeline of behavioral assessments following self-administration and schematic of operant set-shifting paradigm depicting levers (gray), dipper (white), cue location (yellow), and correct response (star). **b-e,** Testing following short-term self-administration. **b,** For number of days to pass lever training criterion (2 consecutive days ≤ 5 omissions), no main effect of sex, treatment, or a sex by treatment interaction was detected (*sex:* F_(1,24)_=1.24, p=0.28; *treatment:* F_(1,24)_=4.09, p=0.054; *interaction:* F_(1,24)_=0.96, p=0.34). **c,** Comparison of trials to criterion for visual cue test showed no significant effect of sex, treatment, or interaction (*sex*: F_(1,24)_=1.07, p=0.31; *treatment:* F_(1,24)_=0.80, p=0.38; *interaction:* F_(1,24)_=1.21, p=0.28). **d,** Comparison of trials to criterion during the extradimensional (ED) shift test showed a significant interaction (F_(1,24)_=9.62, p=0.005), with remifentanil females requiring more trials versus saline females (p=0.002) and remifentanil males (p=0.002), whereas performance in remifentanil males did not differ compared to saline males (p=0.33). **e,** Comparison of trials to criterion during a reversal test showed no significant main effects or interaction (*sex*: (F_(1,19)_=0.05, p=0.82); *treatment:* (F_(1,19)_= 0.12, p=0.73); *interaction:* (F_(1,19)_=0.24, p=0.63). **f-h,** Testing following long-term self-administration in males. **f-g**, No difference was observed in number of days to pass lever training criterion (t_(13)_=−0.54, p=0.60) or trials to reach criterion during a visual cue test (U=18.00, p=0.28). **h,** During the ED test, remifentanil males required more trials to reach criterion compared to saline (t_(13)_=−2.64, p=0.02). *p<0.05, **p<0.01.

As a similar hypoactive basal state was observed in males following long-term self-administration, we next measured cognitive flexibility in males following prolonged remifentanil exposure. No differences in days to reach lever training criterion or trials to reach criterion during the visual cue test were observed (Figure 5f-g). However, similar to females following short-term self-administration, remifentanil males required significantly more trials to reach criterion during the extradimensional shift test (Figure 5h). These findings indicate that opioid self-administration results in cognitive flexibility deficits in both males and females, and that this deficit arises in females with less drug exposure and aligns with emergence of a hypoactive basal state.

### Chemogenetic manipulation of prelimbic function and cognitive flexibility

If the emergence of a hypoactive basal state underlies reductions in cognitive flexibility following repeated drug use, induction and compensation of this state should impair and restore flexibility, respectively. To test this possibility, we used an *in vivo* chemogenetic approach to express the inhibitory hm4d-Gi designer receptor exclusively activated by designed drugs^56^ (DREADD) in CamKII-expressing cells (i.e. pyramidal neurons) during the extradimensional shift test in females that previously self-administered saline (Figure 6a). Following a habituating injection of saline, there were no differences in trials to criterion during a visual cue test (Figure 6b). Similar to previous prelimbic lesion studies^57^, injection of the DREADD agonist clozapine-N-oxide (CNO; 2.0 mg/kg i.p.; 30 min prior) in mice expressing the Gi-DREADD took significantly more trials to reach criterion during the extradimensional shift test compared to mice with a GFP-control virus (Figure 6c). *Ex vivo* analysis of Gi-DREADD effects on pyramidal neuron excitability following *in vivo* CNO injection showed that hM4Di expressing cells had significantly higher rheobases compared to cells from GFP-expressing mice (Figure 6d).

**Figure 6.**
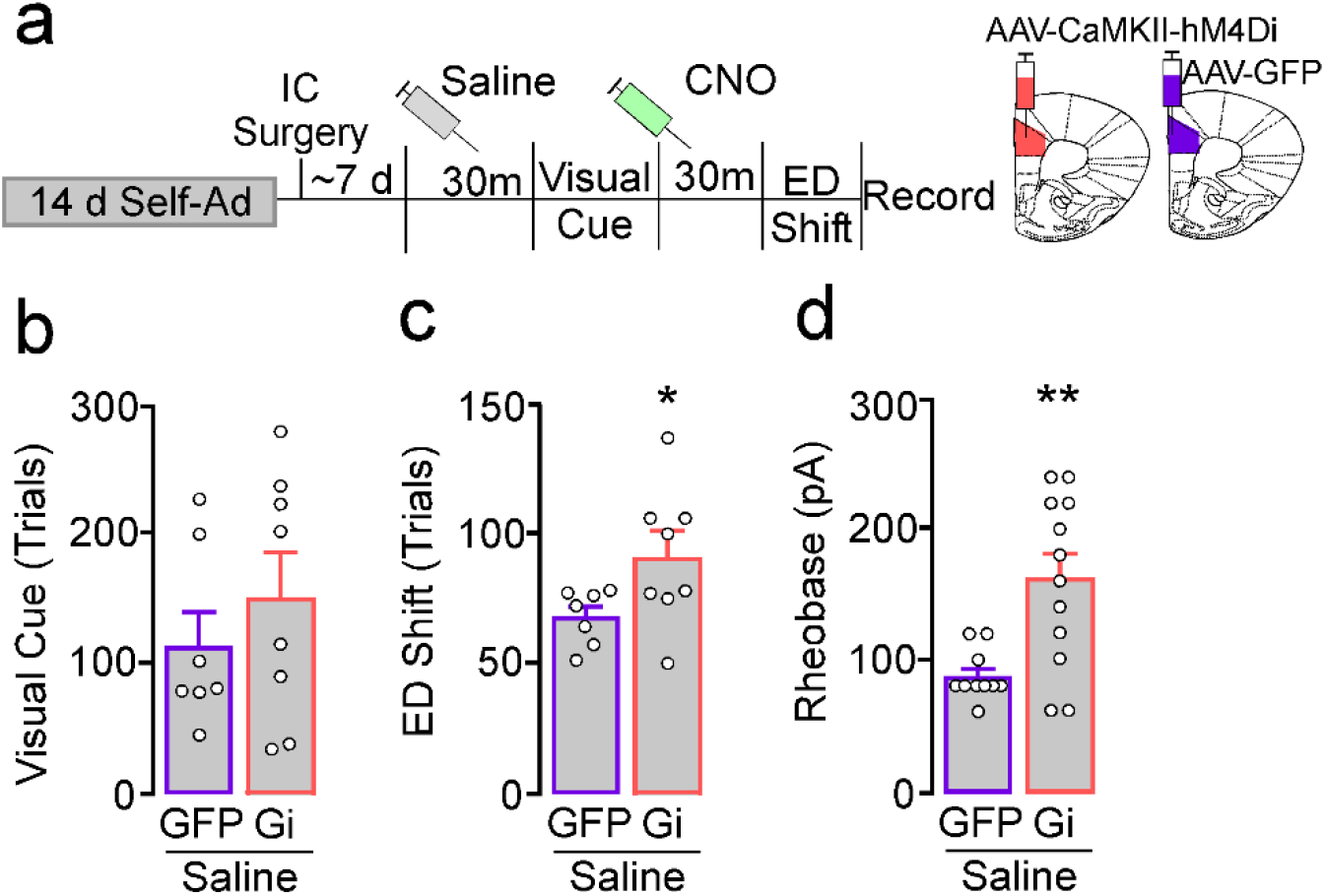
Activation of a prelimbic Gi-DREADD in females induces deficits in cognitive flexibility. **a,** Timeline of DREADD experiments involving short-term saline females expressing a control GFP AAV or a CAMKII-hm4d(Gi)-DREADD in the prelimbic cortex. All mice received saline or Clozapine-N-Oxide (CNO; 2.0mg/kg, i.p.) injection 30 minutes prior to visual cue and ED shift test, respectively. **b,** Comparison of trials to criterion during visual cue testing showed no difference between GFP and Gi-DREADD female mice (t_(13)_ = −0.86, p=0.41). **c,** For trials to criterion during the ED shift, Gi-DREADD females required more trials compared to GFP controls (t_(13)_= −2.16, p=0.05). **d,** *Ex vivo* comparison of rheobase in prelimbic L5/6 pyramidal neurons following systemic CNO injection. Pyramidal neurons expressing Gi-DREADD (N/n=6/12) showed increased rheobase compared to GFP-expressing cells (N/n=5/10; t_(20)_=−3.41, p=0.003). *p<0.05, **p<0.01.

We next used a similar approach to express the excitatory hm3d-Gq DREADD in female mice that underwent either saline or remifentanil self-administration (Figure 7a). As above, no differences were observed following a saline injection during the visual cue test (Figure 7b). Following a CNO injection, GFP-expressing remifentanil females took significantly more trials to reach criteria compared to GFP-saline during the extradimensional shift test, replicating our initial findings. Gq-expressing remifentanil females performed significantly better than GFP-expressing remifentanil female mice, while performance in Gq-saline mice was similar to their GFP counterparts (Figure 7c). Notably, initial studies using a higher dose of CNO (5 mg/kg) produced a similar rescue of behavior in Gq-expressing remifentanil females; however, it also rescued performance in GFP-expressing remifentanil female mice when compared to our initial dataset (Supplemental 10), suggesting at higher concentrations, back metabolism of CNO to clozapine^58^, which improves set-shifting performance in rats^59^, has off-target effects.

**Figure 7.**
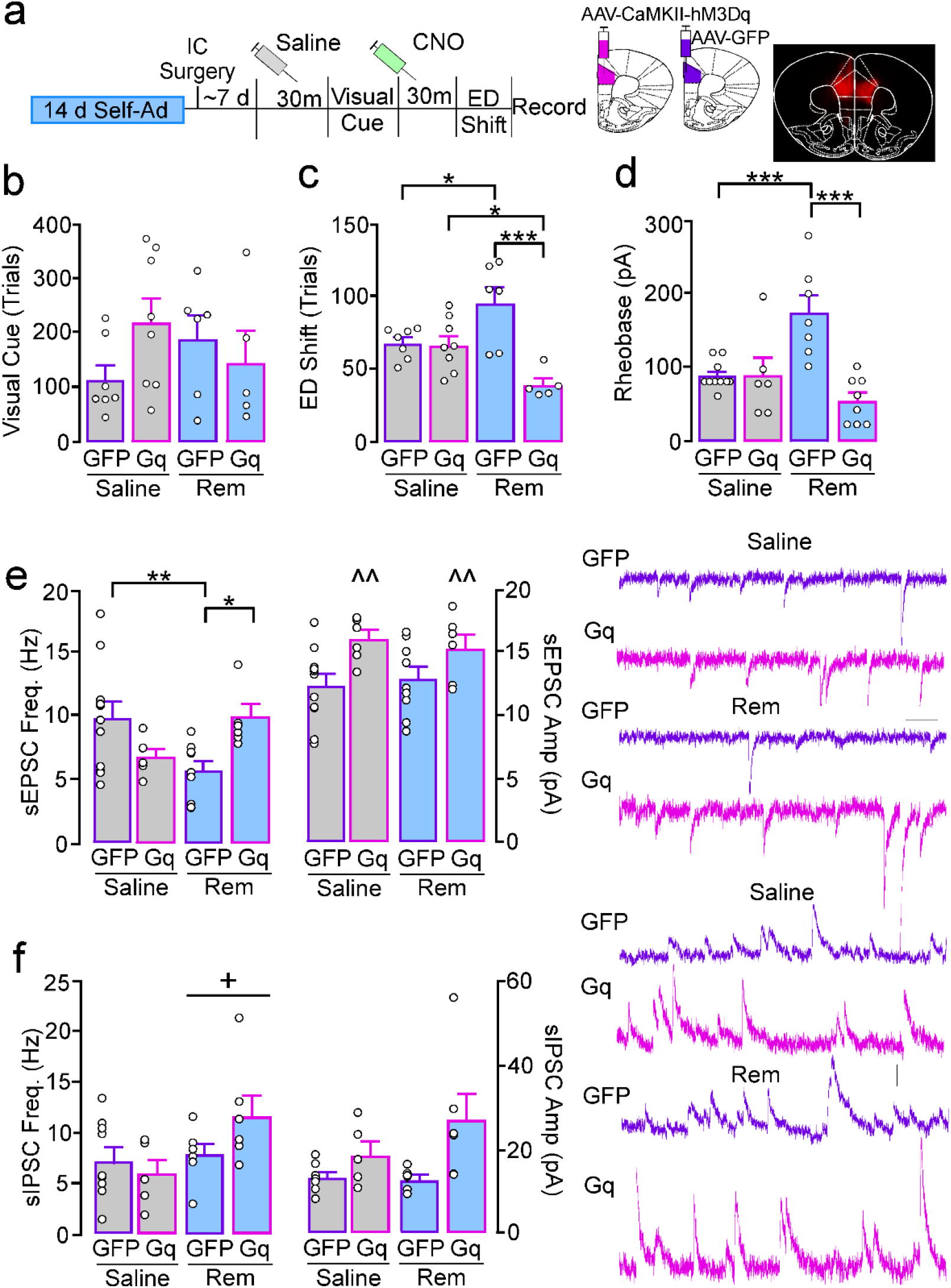
Increased excitation of prelimbic pyramidal neurons in remifentanil females reverses deficits in cognitive flexibility. **a,** Timeline of DREADD experiments involving short-term saline and remifentanil female mice expressing an excitatory CaMKII-hM3Dq(Gq)-DREADD or control GFP AAV in the prelimbic cortex. All mice received a saline or CNO (1.5 mg/kg, i.p.) injection 30 minutes prior to the visual cue and ED shift test, respectively. **b,** Comparison of trials to criterion during visual cue testing showed no effect of treatment, virus, or treatment by virus interaction (*treatment:* F_(1,22)_= 0.00, p=0.97; *virus:* F_(1,22)_= 0.55, p=0.46; *interaction:* F_(1,22)_= 2.86, p<0.11). **c,** For trials to criterion during the ED shift, a significant treatment by virus interaction was observed (F_(1,22)_= 13.52, p=0.001). Post-hoc comparisons showed that female GFP-remifentanil mice required more trials versus GFP-saline (p=0.016) and Gq-remifentanil mice (p<0.001). Gq-remifentanil mice performed significantly better than gq-saline mice (p=.017). There was no significant difference between GFP-saline and Gq-saline (p=0.86). **d-f,** *Ex vivo* comparison of rheobase and synaptic transmission following systemic CNO injection. **d,** Rheobase in prelimbic pyramidal neurons showed a treatment by virus interaction (F_(1,27)_= 14.98, p<0.001), with more current required to evoke an action potential in female GFP-remifentanil mice (N/n=3/7) versus GFP-saline (N/n= 5/10; p<0.001) and Gq-remifentanil mice (N/n=4/8; p<0.001). Rheobase in Gq-saline did not differ compared to GFP-saline (N/n= 4/6; p=0.93) or Gq-remifentanil (p=0.12). **e,** Spontaneous EPSC (sEPSC) frequency **(left)** and amplitude **(right)** in GFP- and Gq-expressing prelimbic pyramidal neurons. For sEPSC frequency, a treatment by virus interaction was observed (F_(1,28)_=10.73, p=0.003). Pyramidal neurons in GFP-remifentanil mice (N/n= 3/9) showed a reduction in frequency compared GFP-saline (N/n= 4/11; p=0.006) and Gq-remifentanil mice (N/n= 3/6; p=0.013). Gq-saline (N/n=3/6) was not significantly different compared to GFP-saline (p=0.058) or Gq-remifentanil (p=0.082). For EPSC amplitude, a main effect of virus, but not treatment or virus by treatment interaction was observed (*virus:* F_(1,28)_= 9.81, p=0.004; *treatment:* F_(1,28)_= 0.00, p=0.94; *interaction:* F_(1,28)_= 0.36, p=0.56), with Gq+ cells exhibiting an overall greater amplitude compared to GFP. **f,** Spontaneous IPSC (sIPSC) frequency and amplitude in GFP and Gq-expressing prelimbic pyramidal neurons. For sIPSC frequency, a main effect of treatment, but not virus or virus by treatment interaction was detected (*treatment:* F_(1,23)_= 4.34, p=0.049; *virus:* F_(1,28)_= 0.55, p=0.47; *interaction:* F_(1,23)_= 2.77, p=0.11). mIPSC frequency was increased in remifentanil mice (GFP:2/7; Gq: 3/6) compared to saline mice (GFP:5/9; Gq:3/5), regardless of virus. For sIPSC amplitude, a Kruskal-Wallis indicated a significant difference between groups (H(3)=9.73, p=0.02) however no pairwise comparisons of interest were significant. Scale bars: 10pA/100msec. *p<0.05, **p<0.01, ***p<0.001, ^^p<0.01 main effect of virus, +p<0.05 main effect of treatment.

**Figure 8.**
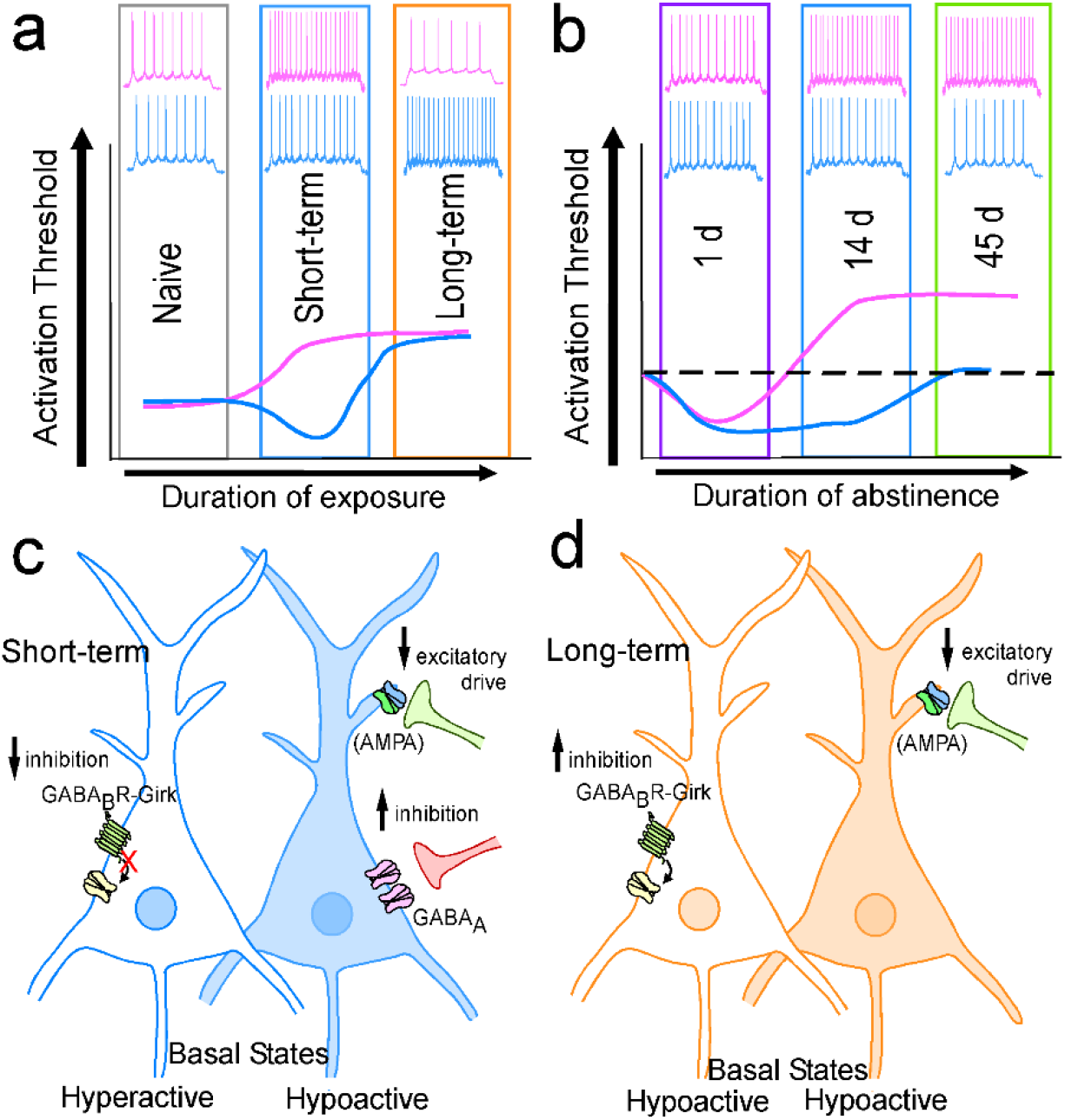
Time- and sex-dependent adaptations in prelimbic pyramidal neuron membrane excitability and synaptic modifications. **a,** Although similar at a naïve state (gray bar), the current series of experiments show that following short-term opioid self-administration (blue bar), L5/6 pyramidal neurons have decreased activation threshold and increased firing in males (blue) but increased threshold firing in females (pink). Following long-term self-administration (orange bar), both males and females show increased activation threshold, however females show reduced firing whereas males show increased firing similar to that following short-term administration in females. **b,** After short-term opioid self-administration, both males and females show reduced threshold to fire an action potential at 1 day abstinence (purple bar), a timepoint where females also show increased firing. At 14-21 days abstinence (blue bar) males show a decreased threshold whereas females show an increased threshold and firing. At 45 days abstinence (green bar), males have returned to naïve levels (dotted black line) whereas females maintain an increased threshold to fire an action potential. **c,** Following short-term self-administration and 14-21 days of abstinence, males (left, unfilled neuron) show a reduction in GABA_B_-GIRK signaling which aligns with a decrease in threshold to fire an action potential. Conversely, females (right, filled neuron) show a reduction in AMPA signaling and an increased in GABA_A_ signaling, both aligning with increased threshold to fire an action potential. **d,** After long-term opioid self-administration and 14-21 days of abstinence, males (left, unfilled neuron) shown an increase in GABA_B_-GIRK signaling which aligns with a hypoactive basal state. Females (left, filled neuron) also show a hypoactive basal state however appears to be driven solely by reductions in AMPA signaling.

*Ex vivo* analysis of CNO effects on pyramidal neuron rheobase showed that CNO bath application (5μM for 10 min) lead to spontaneous firing in 4 out of 5 cells and that CNO also significantly reduced rheobase in non-virus expressing and GFP-expressing cells, albeit to a lesser extent (Supplemental 10). *Ex vivo* physiological assessments following *in vivo* systemic injection of CNO showed that, similar to findings in non virus-expressing animals, rheobase in GFP-expressing remifentanil female mice was significantly greater than GFP-expressing saline mice. Alternatively, rheobase in Gq-remifentanil mice was significantly reduced compared to cells from GFP-remifentanil and similar to GFP- and Gq-saline (Figure 7d). Assessment of synaptic transmission following *in vivo* CNO showed that sEPSC frequency was reduced in GFP-remifentanil mice compared to GFP-saline controls and Gq-remifentanil mice (Figure 7e). Additionally, CNO in Gq mice produced an overall increase in sEPSC amplitude regardless of treatment (Figure 7e). Assessment of sIPSCs showed that remifentanil mice, regardless of virus, had greater sIPSC frequency compared to cells from saline mice, with no differences in sIPSC amplitude (Figure 7f). Taken together, these data demonstrate that opioid-induced hypoactive basal states underlie impairments in cognitive flexibility, and that this hypoactive state is ostensibly driven by a reduction in excitatory drive at pyramidal neurons.

## Discussion

Drug-induced dysfunction of essential cognitive functions strengthen drug-related behavior, increase the risk of developing out of control drug use, and hinder sustaining abstinence^7,11–15^. Females are at heightened risk for developing substance abuse disorders, and exposure to illicit substances leads females to transition to addiction more rapidly than men^60,61^. While opioid-induced adaptations in male and female mice cannot definitively be mapped on to alterations in humans, we outline a potential mechanism by which opioids promote a transition to addiction, and why this occurs more rapidly in females. We demonstrate that low dose exposure of a clinically used opioid is sufficient to promote an enduring hypoactive basal state in prelimbic pyramidal neurons in male and female mice akin to reduced basal activity observed in individuals addicted to cocaine and heroin^7,16^. This dysfunction occurs on a faster timeline in females and aligns with the emergence of deficits in cognitive flexibility commonly observed in human addicts^7,11–15^. A striking feature of this study is that these hypoactive states are driven by distinct cellular mechanisms, reflecting increased inhibition mediated by GABA_B_R but not GABA_A_R in males and reduced excitatory (AMPAR) drive in females.

### Time-dependent alterations in pyramidal neuron function

Abstinent heroin and cocaine users show reductions in basal metabolic activity brain regions homologous to the prelimbic that coincide with craving related increases in neural activity in response to drug cues^24–28^. A similar phenomenon has been observed in separate rodent cocaine studies with males, showing hyperexcitablity of prelimbic pyramidal neurons and increased responsivity during early cocaine use^31,39,40,62,63^ and a hypoexcitable basal state following eight weeks of self-administration^29^. This dichotomy aligns with our current findings, where prelimbic pyramidal neurons exhibit increased rheobase but also increased firing frequency at more depolarized currents. This phenomenon is observed following 14 days of self-administration in females but takes 30 days in males. Increased action potential firing, along with increased variability, appears to be driven by a small portion of cells which may reflect pathway-specific adaptations, whereby increased firing capacity is confined to circuits that drive drug-seeking such as prelimbic projections to the nucleus accumbens core^18,64–67^. Alternatively, the increased firing and variability could reflect time-specific plasticity, as this was not observed following long-term remifentanil self-administration in females, nor was it observed in males self-administering cocaine for 8 weeks^29^. As PFC hypofunction is thought to coincide with strengthening of habit-related circuits, the lack of heightened firing following long-term exposure in females may reflect a devolution in behavioral control from prefrontal to dorsal striatal circuits^7,13,29,68–76^ that is not yet evident in males after only 4 weeks of exposure.

A wide body of work has identified time-dependent adaptations in prelimbic activation states and plasticity-related genes during drug withdrawal^31,42–45^. We find that the increased rheobase in females but not the decreased rheobase in males lasts up to at least 45 days, suggesting that this adaptation may contribute to enduring deficits in behavioral control. Interestingly, females showed a hyperactive profile characterized by reduced rheobase and increased action potential firing during acute withdrawal (24-72 hours). The functional implication of this shift is unclear but may represent a permissive functional plasticity^77^, whereby lowering the threshold of further synaptic changes to develop.

Unlike cocaine, where the prelimbic and infralimbic play opposing roles in drug-seeking, activation of both sub regions is necessary for heroin seeking^18,19,33,65,78–81^. It was therefore unexpected that exposure to neither short- or long-term remifentanil self-administration resulted in infralimbic changes of basal state or action potential firing in males or females. Consistent with our array of prelimbic changes following varying lengths of abstinence (changes at 24-72h versus 14-21 days), changes in the infralimbic following opioid self-administration may be time-dependent. However, the variability in rheobase and spike firing across cells more likely suggests that evident changes are occurring in a cell-type specific (i.e., neurons expressing D1- versus D2-type receptors)^82^ or pathway-specific manner, as past work has identified opioid-induced plasticity at infralimbic-inputs to the nucleus accumbens shell^83^.

### Sex-specific excitatory and inhibitory plasticity

Imbalances in cortical excitatory and inhibitory signaling have been implicated in the pathophysiology of impaired cognition associated with numerous disorders^20–22,35,84–87^. Previous studies have implicated mPFC excitatory and inhibitory synaptic modifications in opioid sensitivity and reinstatement^28,88–91^; however, these studies did not distinguish sub regions of the mPFC and assessed plasticity following extinction^35–38^. Although previous work in male rats demonstrated that long-term cocaine administration reduces excitability of prelimbic pyramidal neurons, an underlying mechanism remained elusive. Here we find that changes in basal state (i.e. rheobase) in males aligns with divergent changes in GABA_B_R-dependent currents. The observed reduction in GABA_B_R signaling is similar to our past findings following repeated cocaine^41^. In both instances, increased excitability was observed within 24 h of the final drug exposure. However, adaptations with non-contingent cocaine emerged following 5 daily exposures, whereas increased excitability in the current study was present following 14 but not 5 days of self-administration. The functional implications of reduced GABA_B_R signaling and increased excitability following short-term exposure are unclear but may be a driving factor in drug-seeking or other drug-related behaviors, as viral-mediated reductions in prelimbic GABA_B_R-GIRK signaling pre-sensitizes male mice to cocaine^41^ and systemic administration of baclofen reduces heroin-induced reinstatement following a similar duration of self-administration in males^92^.

In females, alterations in GABA_B_R currents were not detected following short- or long-term exposure. Following short-term exposure, pyramidal neurons in females showed a bidirectional increase and decrease in GABA_A_R- and AMPAR-mediated currents, respectively. However, with more prolonged exposure, only alterations in AMPAR signaling remained, indicating that reductions in excitatory drive are the primary mechanism driving hypoactive basal states and cognitive dysfunction. In support, *ex vivo* assessment following *in vivo* CNO injection showed that even though cognitive deficits and hypoactive basal states remained, CNO abolished increases in mIPSCs in GFP-remifentanil female mice previously observed following short-term self-administration in non-viral cohorts. Further, restoration of cognitive performance and drug-induced hypoactive states in Gq-remifentanil female mice aligned with restoration of (increased) excitatory transmission to control levels, whereas inhibitory transmission was in fact elevated to levels observed in our initial remifentanil cohorts, suggesting that elevations in GABA_A_R signaling are insufficient to drive cognitive decline. These findings are in agreement with past work demonstrating that alterations in excitatory signaling are more disruptive to cortical information flow and mPFC-dependent behaviors than increased cortical inhibition^35^. It is possible that augmentation in inhibitory signaling contributes more to drug-related behaviors, as increased responsivity of mPFC GABAergic interneurons to heroin-associated stimuli has been previously implicated in relapse vulnerability^91^. Notably, the lack of effect on sIPSCs in GFP-remifentanil female mice does not likely reflect a lack of replication but rather a distinction in effects on activity-dependent versus quantal transmission or that back metabolism of CNO alters GABA_A_R signaling. Interestingly, rescue of deficits during the extradimensional shift in Gq-remifentanil mice aligns with a significant reduction in rheobase compared to GFP-remifentanil mice, such that it was similar to GFP-saline mice. No impact of Gq-DREADD on rheobase in saline controls was observed, however this lack of effect aligns with a lack of behavioral change. Further, it likely reflects complexity of cortical microcircuits and functionality of mechanisms normally in place to counter acute shifts in cortical activation.

### Impact of opioid exposure on elevated plus maze and forced swim test behaviors

The PFC regulates systems that drive affect and reward-related behavior^93–97^. The lack of observed changes in the EPM may reflect the use of a short-lasting opioid or the low dose of remifentanil. In preclinical models, somatic withdrawal symptoms are often precipitated using the mu opioid receptor antagonist, naloxone, and/or use significantly higher dependence-inducing doses^98–100^. Our timeline of testing may have also impacted the detection of behavioral changes, specifically in the elevated plus maze, which was done 24-72 hours following the completion of self-administration, a time point we show reduced rheobase in both males and females.

## Conclusion

Our capacity to prevent and treat opioid use disorders is hindered by variability within diagnosed populations. Biological sex is known to dictate drug-related behavioral and neurobiological outcomes^61,101,102^. While not all opioid-related behaviors vary across sex^103,104^, recent work has demonstrated a role of estrous stage on cue/drug associations ^105^, heroin self-administration^106^, and unconstrained remifentanil demand^107^. Thus, a critical step in future studies will be determining what role, if any, gonadal hormones have on the observed deficits in flexibility and how these deficits confer susceptibility to out of control drug use. Further, while opioid addiction appears to share a number of core pathologies with addiction to other drugs of abuse^18,66,67,81,108^, literature also support the contention that addiction to opioids exhibits behavioral and neurobiological distinctions^109–112^. Thus, while information gained from previous work with non-opioid drugs has laid the groundwork for understanding addiction pathology, we cannot assume that the same mechanisms underlie emergence of opioid use disorders. Here, we have identified sex-specific and time-dependent plasticity within a key substrate for cognitive control following opioid exposure. These data highlight important considerations for clinical opioid use, as they show females are at greater risk for PFC dysfunction and impaired behavioral control on a shorter timescale than previously thought. Further, they indicate that interventions aimed at preventing and treating opioid use disorder must be tailored based on biological sex and the neuroadaptations that emerge during different phases of the addiction cycle.

## Acknowledgements

These studies were supported by funding from the National Institute on Drug Abuse grant K99 DA038706 (M.H.), and R00DA038706 (M.H.), as well as funding from the Brain and Behavior Research Foundation (#26299).

---

**Supplemental Figure 1.**
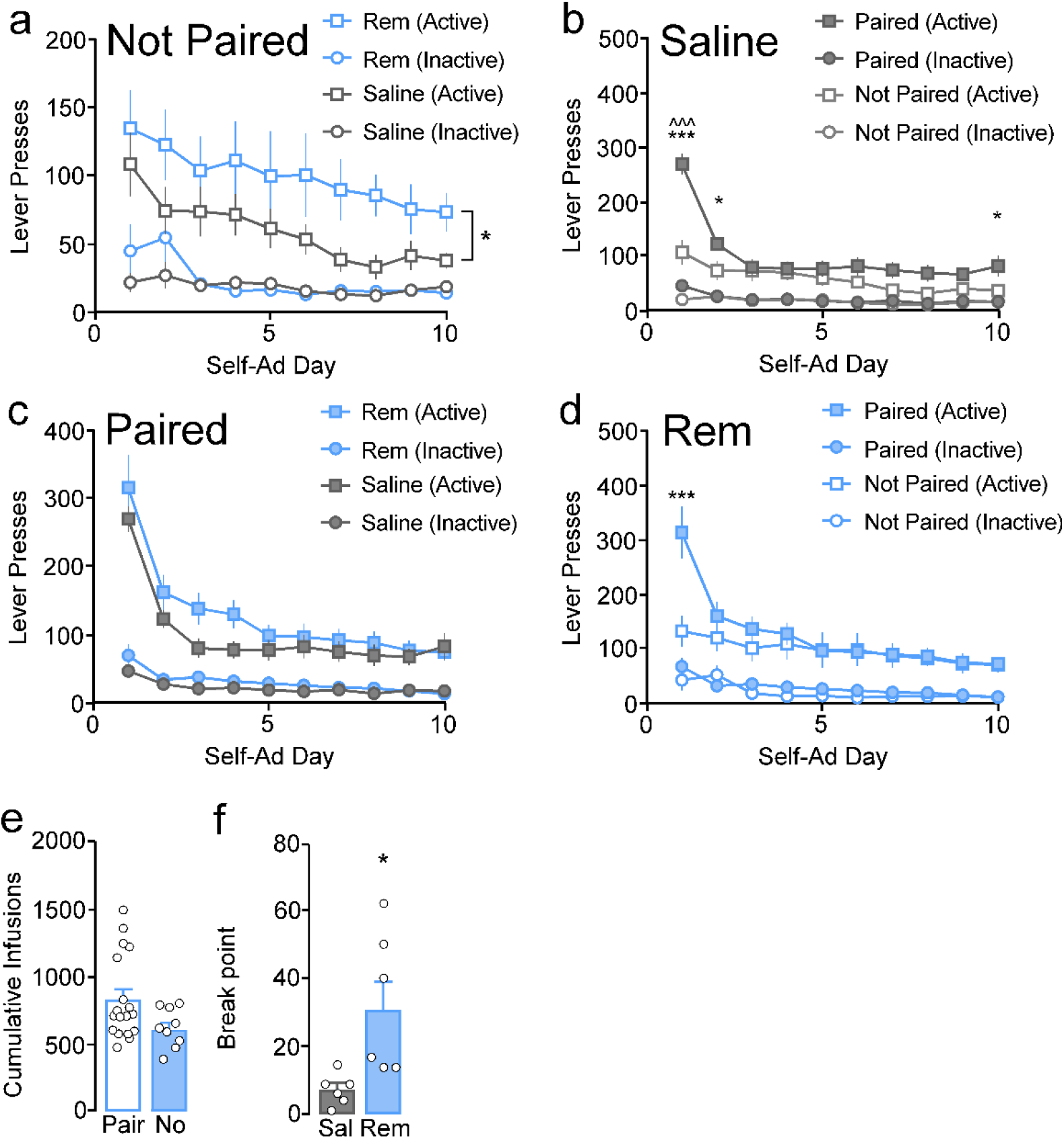
Paired Ensure®-Infusion training increases active lever pressing in saline, but not remifentanil, male mice. A portion of mice were trained using Ensure® paired with remifentanil or saline for three days on an increasing fixed ratio schedule (paired) whereas others received a single day of Ensure® that was not paired with an infusion (not paired). **a,** When ensure was not paired with an infusion during training, remifentanil mice had overall greater active lever presses compared to saline mice (F_(1, 27)_= 4.60, p=0.04) and there was a main effect of day (F_(9,243)_= 4.69, p<0.001), however there was no treatment by day interaction (F_(9,243)_= 0.19, p=0.995). For inactive presses, there was a treatment by day interaction (F_(9,243)_= 1.96, p=0.045), with remifentanil mice having greater inactive presses on day 1 (p=0.02) and 2 (p=0.006). **b,** When Ensure® was paired with the infusion using an increasing fixed ratio schedule during acquisition, saline and remifentanil mice had similar active and inactive lever presses during the first ten days of self-administration (*active*: F_(1,32)_= 1.57, p=0.22; *inactive:* F_(1,32)_= 2.29, p=0.14) and there was no treatment by day interaction (*active*: F_(9,288)_=1.05, p=0.40; *inactive*: F_(9,288)_= 1.02, p=0.42), however overall there was a decrease in presses across days (*active*: F_(9,288)_ = 40.74, p<0.001; *inactive*: F_(9,288)_=10.21, p<0.001). **c,** Paired male mice receiving saline infusions prior to saline self-administration had significantly more active lever presses (gray filled, square) than saline Unpaired males (interaction: F_(9, 306)_= 7.97, p<0.001, training type: F_(1, 34)_= 7.42, p=0.01). Paired training mice decreased inactive presses (circles) over days to a greater extent than not paired mice (interaction: F_(9,306)_= 2.07, p=0.03, training type: F_(1, 34)_= 0.53, p=0.47). **d,** There was a day by training interaction on active lever presses (F_(9, 225)_= 4.18, p<0.001) however higher active lever responding was only observed in paired versus not paired remifentanil mice on the first day following Ensure® training (p<0.001). Paired mice had similar inactive lever pressing compared to not paired mice (*interaction*: F_(9, 225)_= 1.23, p=0.28, *training type*: F_(1, 25)_= 1.53, p=0.23). **e,** Paired mice had similar total infusions compared to unpaired mice (t_(25)_=−1.95, p=0.06) and total infusions did not correlate to average prelimbic rheobase for not paired (p=0.88) or paired (p=0.73) mice. **f,** A subset of saline and remifentanil male mice that were trained to lever press using the paired (i.e. ensure paired with infusion) went through a progressive ratio test on the last day of self-administration. Remifentanil mice had a significantly higher breaking point compared to saline mice (U= 5.00, p=0.04). ***p<0.001 paired vs not paired active presses, *p<0.05 paired vs not paired active, ^p<0.05 paired vs not paired inactive.

**Supplemental Figure 2.**
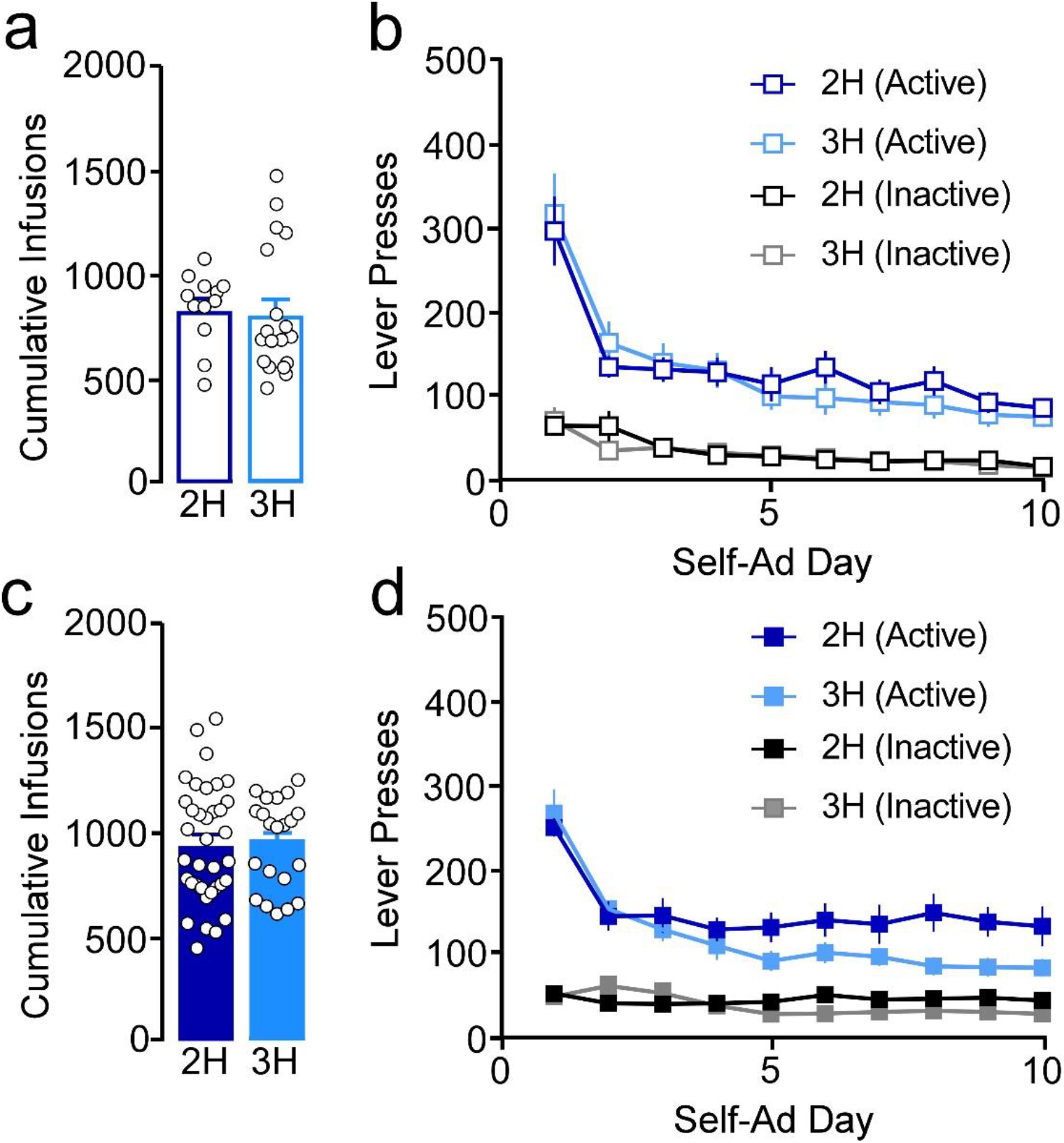
Duration of self-administration session does not contribute to overall remifentanil infusions earned. In order to determine if duration of self-administration contributed to overall responding and intake (infusions), male and female mice were run for 2- or 3-hour daily self-administration sessions. **a,** Mean total infusions in 2 hour and 3 hour self-administering males show no significant difference (t_(28)_=0.23, p=0.82). **b,** In males, there was no difference in active (*session length*: F_(1,28)_= 0.06, p=0.81; *interaction*: F_(9,251)_= 0.65, p=0.75) or inactive lever presses (*session length*: F_(1,28)_= 0.15, p=0.70; *interaction:* F_(9,251)_= 0.97, p=0.46). **c,** Similar to males, no differences in total remifentanil intake was observed between 2 and 3 hour females (t_(55)_= −0.12, p=0.90). **d,** In females, there was a session length by day interaction for active (F_(9,513)_= 2.30, p=0.02) and inactive lever presses (F_(9, 513)_=3.40, p<0.001). The only significant post-hoc comparison was that 2 hour females had significantly more active presses on day 8 compared to 3 hour females (p=0.024).

**Supplemental Figure 3.**
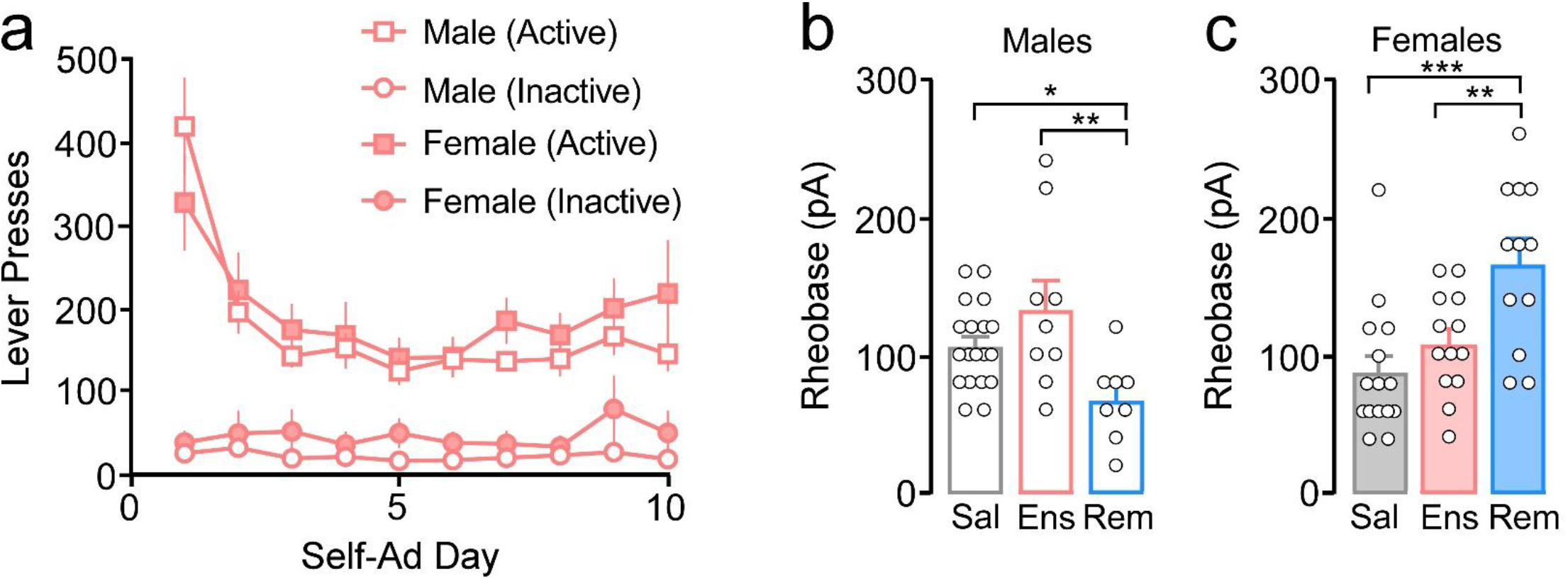
Ensure® self-administration does not alter L5/6 pyramidal neuron membrane excitability in the prelimbic cortex. **a,** There was no main effect of sex or a sex by day interaction on active (*sex:* F_(1,15)_= 0.29, p=0.60; *interaction:* F_(9,135)_= 1.61, p=0.12) or inactive (*sex:* F_(1,15)_= 1.59, p=0.23; *interaction:* F_(9,135)_= 1.06, p=0.40) presses. **b,** In males, there was a significant effect of treatment on rheobase (F_(2,33)_= 5.87, p=0.007). Post-hoc comparisons show remifentanil (rem) mice (N/n= 6/8) have significantly lower mean rheobase compared to saline (sal) mice (N/n=13/19; p=0.027) and ensure (ens) mice (N/n=3/9; p=0.005), with no difference between ensure and saline (p=0.10). **c,** In females, there was a significant effect of treatment on rheobase (F_(2,37)_= 9.30, p<0.001). Post-hoc comparisons show remifentanil mice (N/n= 4/12) have significantly higher mean rheobase compared to saline (N/n= 9/15; p<0.001) and ensure (N/n= 5/13; p=0.005), however no difference between ensure and saline (p=0.25). ***p<0.001; **p<0.01; *p<0.05.

**Supplemental Figure 4.**
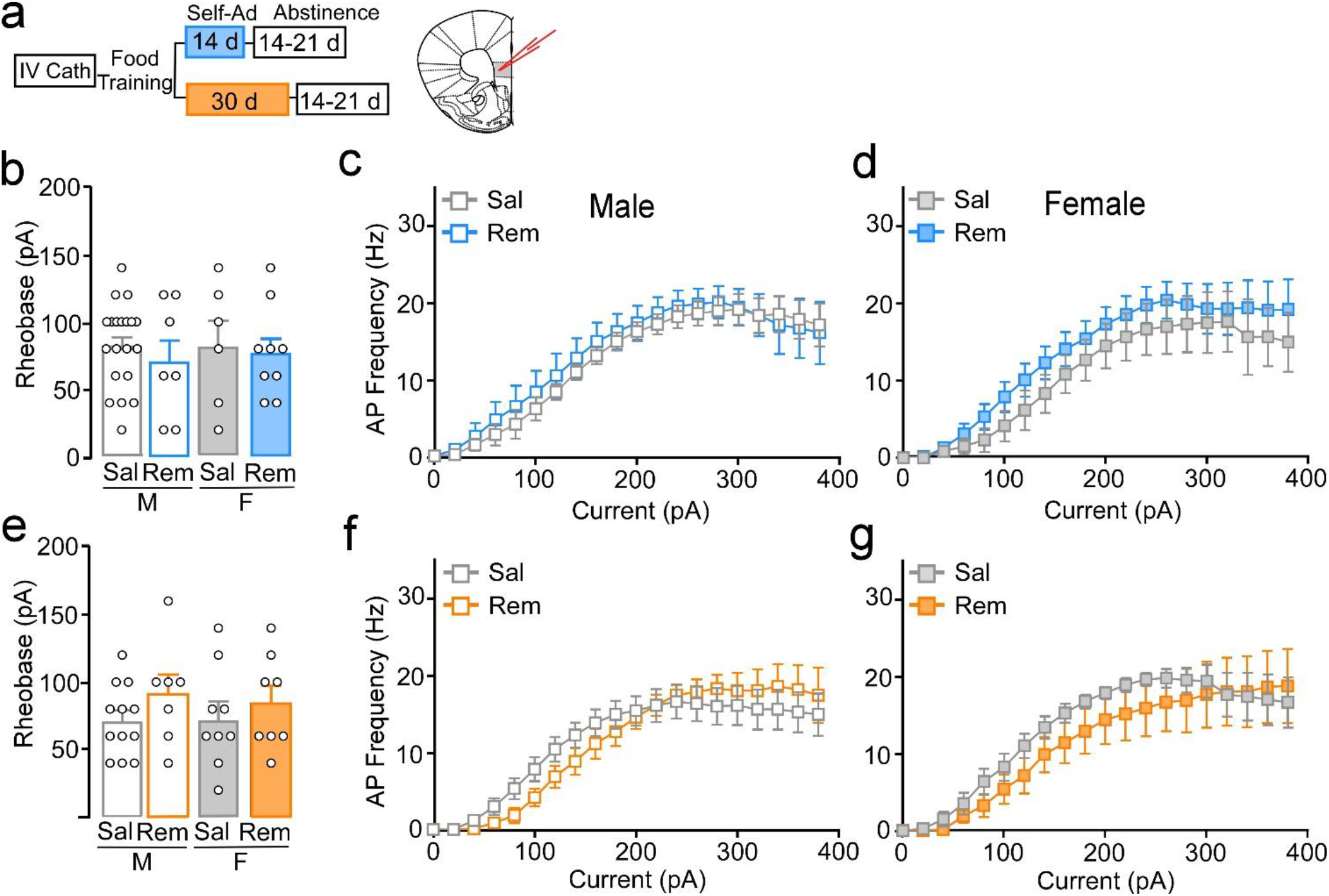
Remifentanil self-administration does not alter L5/6 pyramidal neuron membrane excitability in the infralimbic cortex. **a,** Self-administration and abstinence timeline and location of current-injection whole-cell recordings, in the infralimbic cortex. **b,** Mean action potential (AP) threshold (rheobase) in infralimbic L5/6 pyramidal neurons following 14-21 d abstinence from short-term self-administration showed no interaction or main effects of sex or treatment (*interaction*: F_(1,38)_= 0.06, p=0.81; *sex*: F_(1,38)_=0.07, p=0.79; *treatment*: F_(1,38)_=0.48, p=0.49; Sal-M: N/n= 9/20, Rem-M: N/n= 5/7, Sal-F: N/n= 5/6, Rem-F: N/n=5/9). **c-d,** Current-spike analysis showed no differences in mean firing frequency in remifentanil (N/n=5/7) versus saline (N/n=7/15) males (**c,** *interaction*: F_(19,380)_= 0.20, p=1.00; *treatment*: F_(1,20)_= 0.31, p=0.59). Current-spike analysis showed no differences in mean firing frequency in remifentanil (N/n=5/9) versus saline (N/n=5/6) females (**d,** *interaction*: F_(19,247)_= 0.21, p=1.00; treatment: F_(1,13)_= 0.97, p=0.34). **e,** Following long-term remifentanil, infralimbic L5/6 pyramidal neurons exhibited similar firing thresholds in males (N/n=3/7) and females (N/n=3/8) compared to respective saline controls (Male: N/n=6/12, Female: N/n=4/9; *interaction*: F_(1,32)_= 0.13, p=0.73, *sex*: F_(1,32)_=0.04, p=0.84; *treatment*: F_(1,32)_=1.90, p=0.18). **f-g,** Current-spike analysis showed no differences in firing frequency across current in remifentanil males (N/n=3/7) compared to saline (**f,** N/n=6/12; *treatment:* F_(1,19)_= 0.02, p=0.90, *interaction:* F_(19,323)_= 1.25, p=0.22) and no differences in remifentanil females (N/n=3/8) compared to saline (**g,**N/n=4/8; *treatment:* F_(1,14)_= 0.50, p=0.49, *interaction:* F_(19,266)_= 0.61, p=0.90).

**Supplemental Figure 5.**
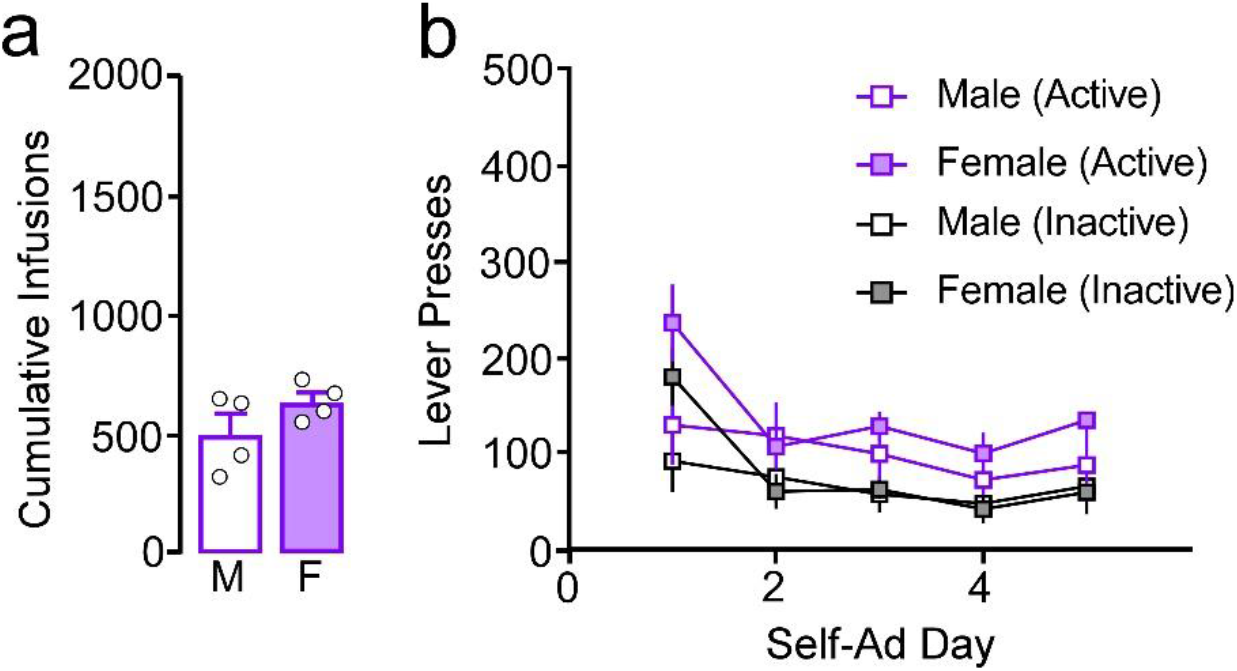
Responding through 5 days of self-administration. **a,** For 5 day groups, no sex by day interaction or main effect of sex was observed for active lever presses [*interaction*: F_(4,24)_= 1.09, p=0.38; *sex*: F_(1,6)_= 3.25, p=0.12)]. No main effect or interaction for inactive lever presses [*interaction*: F_(4,24)_= 1.92, p=0.14; *sex*: F_(1,6)_= 0.68, p=0.44)]. **c,** Cumulative remifentanil infusions did not differ in males (M) versus females (F) (t_(6)_= −1.47, p=0.19).

**Supplemental Figure 6.**
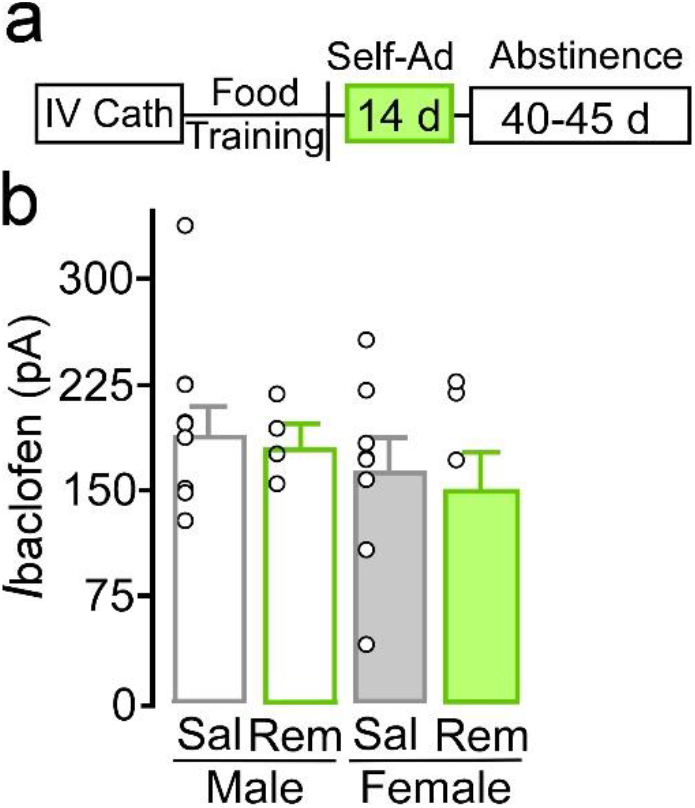
GABA_B_-GIRK signaling changes following remifentanil self-administration are not enduring. **a,** Self-administration and abstinence timeline. **b,** No significant effect of sex, treatment, or interaction on I_Baclof_ en was observed following 40-45 days of abstinence (*sex:* F_(1,24)_= 1.88, p=0.18; *treatment:* F_(1,24)_= 0.24, p=0.63; *interaction:* F_(1,24)_= 0.04, p=0.85).

**Supplementary Figure 7.**
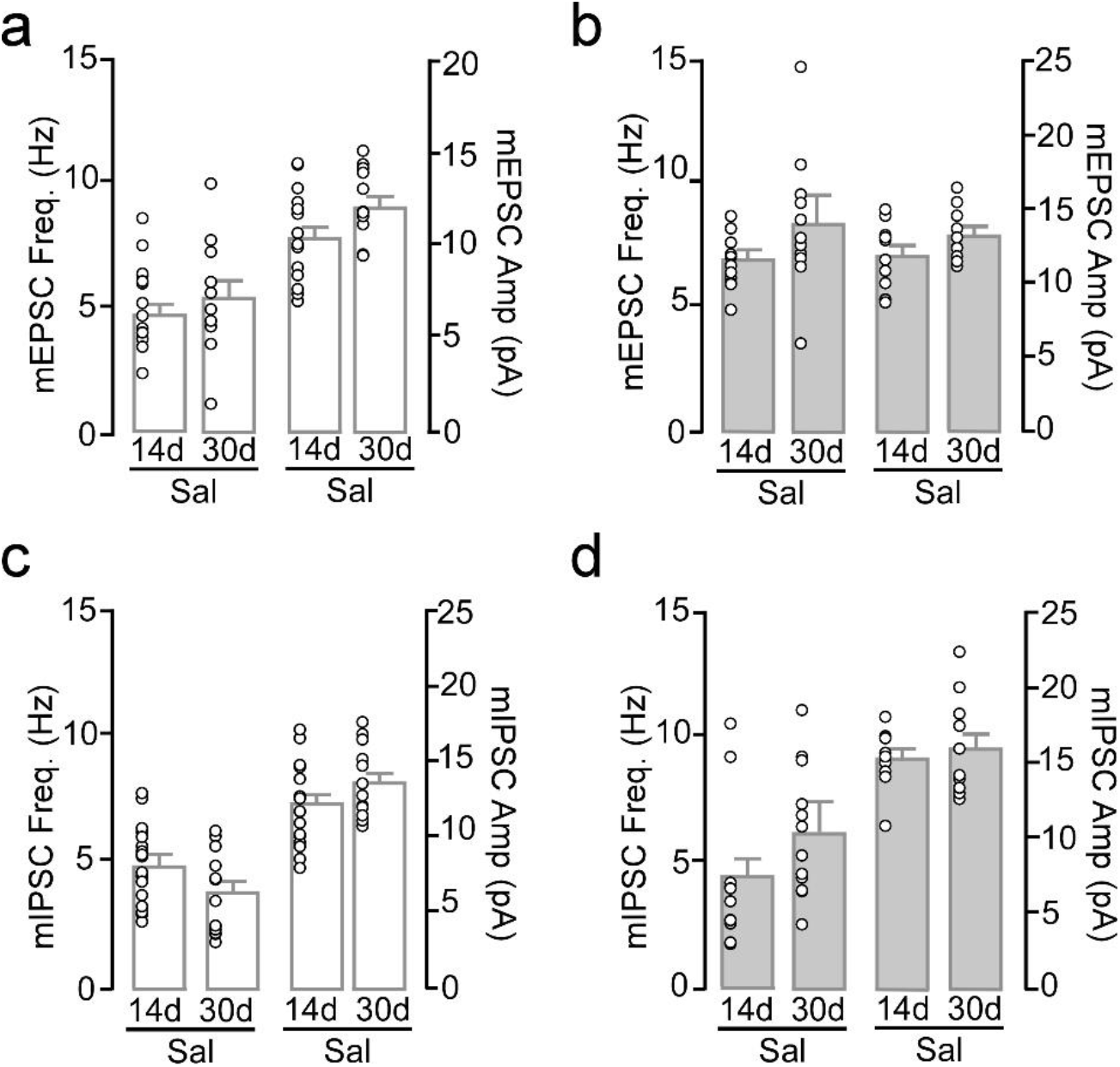
Excitatory and inhibitory transmission in 14 and 30 day saline self-administering mice. **a,** Short- (N/n=8/16) or long-term (N/n=5/11) male saline mice did not differ in mean EPSC frequency (**left**; t_(25)_= −0.93, p=0.36) or amplitude (**right**; t_(25)_= −1.87, p=0.07). **b,** No difference in mean mEPSC frequency (t_(19)_= −1.94, p=0.07) or amplitude (t_(19)_= −1.68, p=0.11) was observed in short- (N/n=6/8) or long-term (N/n=5/13) saline females. **c,** Mean mIPSC frequency (t_(28)_=1.91, p=0.07) and amplitude (t_(28)_=−1.41, p=0.17) did not differ in saline short- (N/n=8/17) versus long-term (N/n=5/13) males. **d,** No difference in mean mIPSC frequency (t_(18)_=−1.77, p=0.09) or amplitude (t_(18)_= −0.58, p=0.57) was observed in short-term (N/n=6/11) versus long-term (N/n=4/9) females.

**Supplemental Figure 8.**
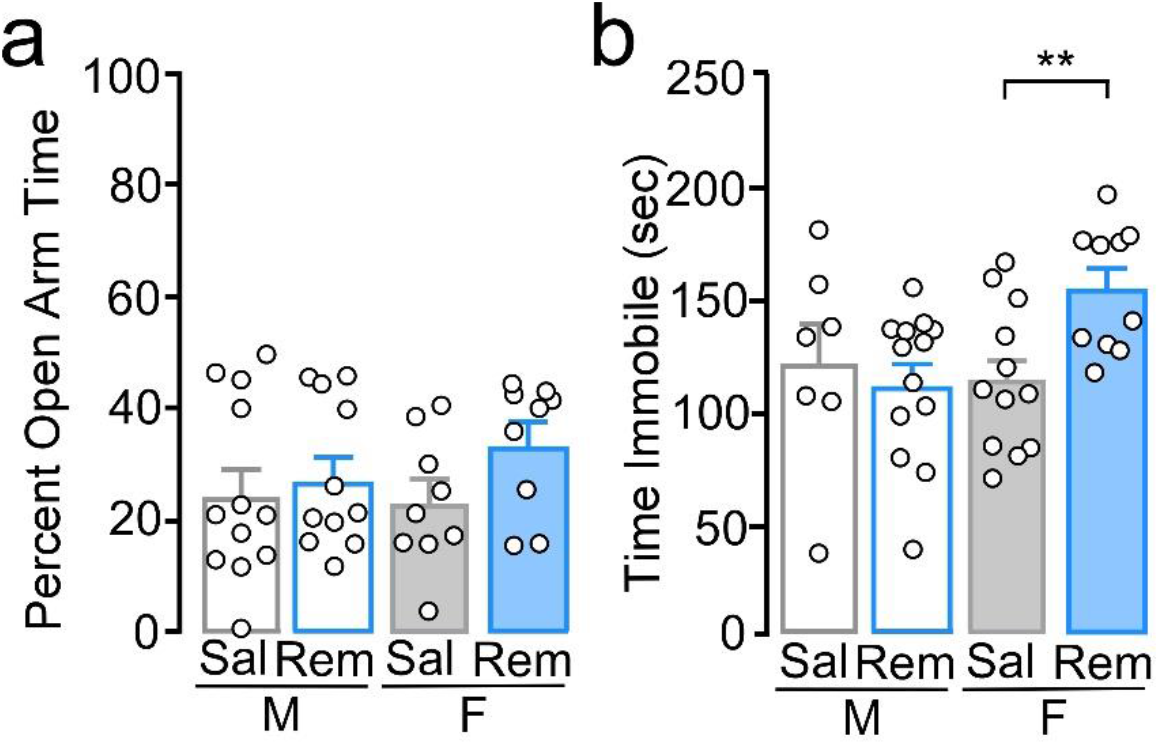
Remifentanil self-administration does not alter percent open arm time in elevated plus maze but increases immobile forced swim time. **a,** Assessment of behavior following short-term self-administration. There was no significant effect of sex (F_(1,37)_= 0.34, p=0.57), treatment (F_(1,37)_= 2.37, p=0.13), or sex by treatment interaction (F_(1,37)_= 0.80, p=0.38) on percent time in the open arm. **b,** There was a significant interaction on time immobile during a forced swim test (F_(1,38)_= 5.56, p=0.02), with remifentanil females spending significantly more time immobile compared to saline females (p=0.007). **p<0.01.

**Supplemental Figure 9.**
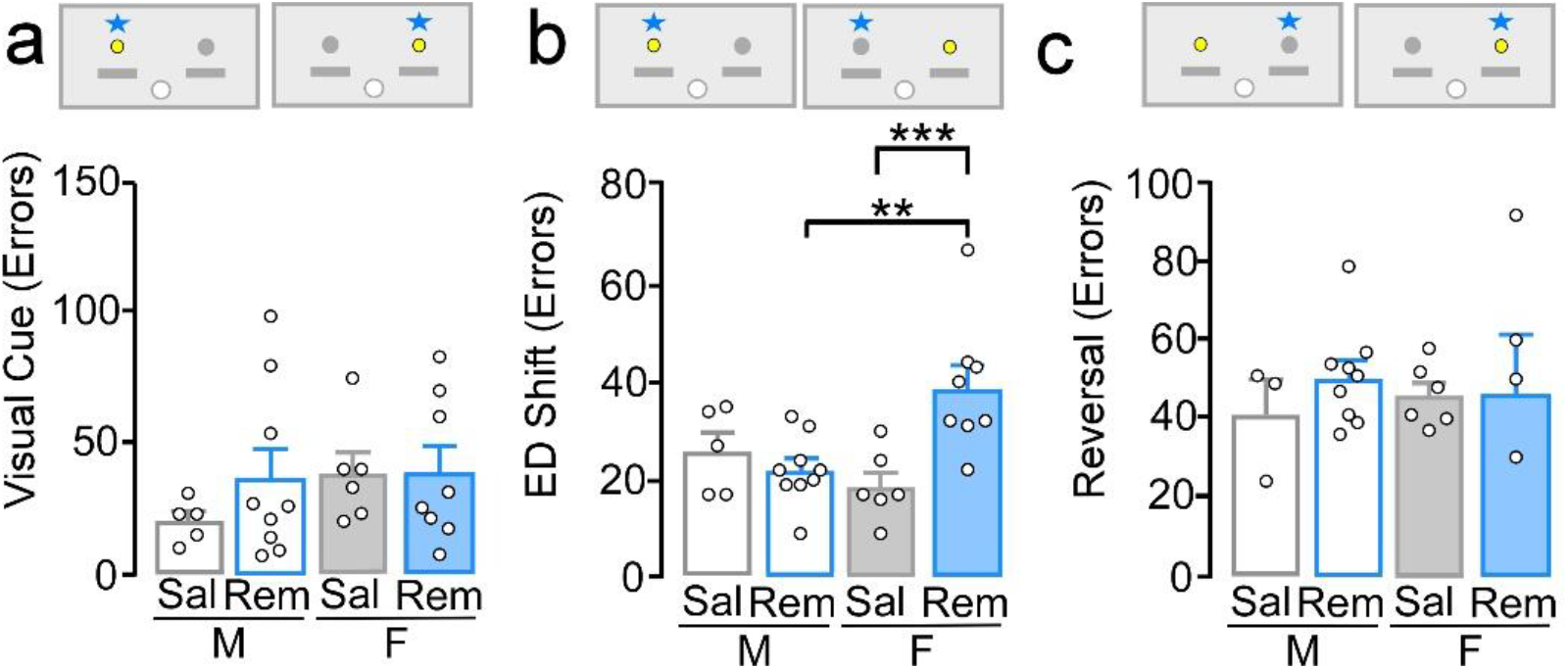
Remifentanil-induced deficits in cognitive flexibility. **a,** Comparison of errors to criterion for visual cue test showed no significant effect of sex, treatment, or interaction (*sex*: F_(1,24)_= 0.97, p=0.34; *treatment:* F_(1,24)_=0.77, p=0.39; *interaction:* F_(1,24)_= 0.65, p=0.43). **b,** Comparison of errors to criterion during the extradimensional (ED) shift test showed a significant interaction (F_(1,24)_= 10.16, p=0.004), with remifentanil females requiring more trials versus saline females (p<0.001) and remifentanil males (p=0.002), whereas performance in remifentanil males did not differ compared to saline males (p=0.48). **c,** Comparison of errors to criterion during a reversal test showed no significant main effects or interaction (*sex*: F_(1,19)_= 0.00, p=0.96; *treatment:* (F_(1,19)_= 0.39, p=0.54; *interaction:* F_(1,19)_= 0.26, p=0.62).

**Supplemental Figure 10.**
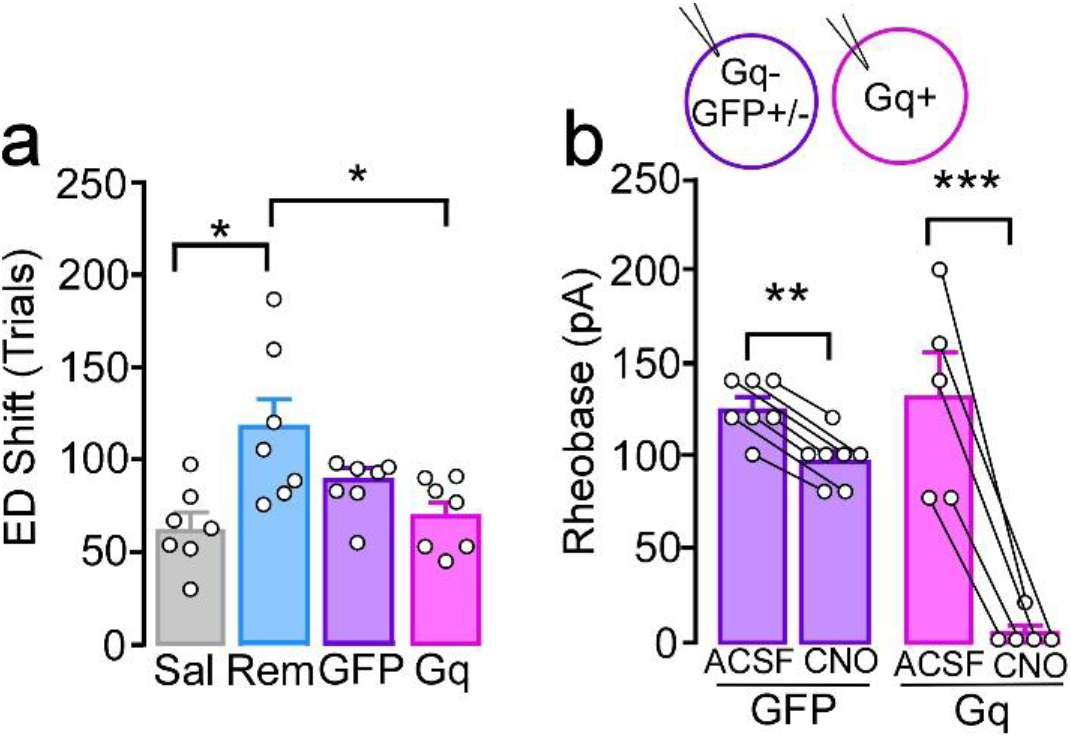
Potential off-target effects of high dose CNO. **a,** During the ED Shift, Gq-remifentanil females (pink, N=7) required fewer trials to criterion versus remifentanil females with no virus (REM, blue; p=0.017), with no difference versus GFP controls (purple, p=0.32) or saline (gray, p=0.66). **b,** *Ex vivo* bath application of CNO (5 μM) reduced rheobase in prelimbic L5/6 pyramidal neurons (F_(1,10)_=78.01, p<0.001), producing spontaneous firing (rheobase = 0) in 4 out of 5 pyramidal neurons positive for Gq-DREADD (N/n= 3/5, p<0.001). Rheobase was also significantly reduced in pyramidal neurons negative for Gq-DREADD and/or positive for GFP-control compared to acsf baseline (N/n=5/7, p=0.032), albeit to a lesser extent (*interaction*: F_(1,10)_= 31.46, p<0.001).

## Online Methods

### Subjects

Adult male and female wild-type C57BL/6 mice (postnatal day 68 ± 0.82 at self-administration onset) were bred in-house or commercially purchased (Jackson Laboratory) and maintained in a temperature and humidity-controlled room. Animal use was approved by the Institutional Animal Care and Use Committee at Marquette University.

### Catheter surgery

Mice were implanted with an intravenous standard mouse jugular vein catheter (Access Technologies; 2/3Fr. x 6cm silicone, Cat No. AT-MJVC-2914A) connected to a backmount (PlasticsOne; 22GA, Part No. 8I31300BM01) under general isofluorane (1-3%) anesthesia. Following catheter surgery, mice were single-housed and allowed to recover for at least 5 days prior to beginning self-administration. During this time, mice were acclimated to handling and catheter manipulation. For all self-administration sessions, catheters were flushed with 0.05 mL (i.v.) of heparinized (20 IU/ml; Hospira, Inc.) bacteriostatic 0.9% saline (Hospira, Inc.) containing gentamicin sulphate (Sparhawk Laboratories, Inc.; 0.25 mg/ml) immediately before and containing enrofloxacin (Norbrook Laboratories; 4.994mg/mL) immediately after each session. Catheter patency was checked periodically using 0.05mL (i.v.) Brevital Sodium (5mg/mL).

### Self-administration

To facilitate acquisition of opioid self-administration, mice were initially food-restricted to maintain a weight of 85-90% their original body weight and were habituated overnight in their home cage to 50% liquid Ensure® (diluted with water). The current study utilizes the highly μOR-specific and potent synthetic opioid, remifentanil HCl (Ultiva®; Mylan Institutional, LLC; purchased from Froedtert Hospital Pharmacy, Milwaukee, WI). During self-administration each intravenous infusion was of either 0.20mL saline or 0.20mL of 5μg/kg/infusion remifentanil based on previous dosing in mice and rats^1–3^.

### ‘Paired’ self-administration paradigm

A variety of self-administration paradigms were used to optimize lever pressing in both male and female mice. ‘Paired’ mice were trained to press the active lever on an increasing fixed ratio schedule using a liquid dipper system (Med Associates, Inc.) paired with an infusion of either saline or remifentanil. Training used a fixed ratio 1 where each active lever press resulted in 20 second presentation of the Ensure® dipper for a total of 25 dipper/infusions. Once all rewards were earned, the following day a fixed ratio 2 was used with a total of 50 dipper/infusion pairings. Lastly, mice had to press the active lever 3 times (fixed ratio 3) for each dipper/infusion presentation and earned a total of 100 reward presentations. The maximum number of reinforcers needed to be earned before proceeding to the next fixed ratio training. Following completion of training, mice were given 1-3 days of abstinence followed by one self-administration session of the remifentanil or saline alone before food was returned *ad libitum.* Following acquisition, male and female mice received remifentanil or saline (average of 5 days per week) on a fixed ratio 1 schedule (+cue-light, time out 20 s) for 2 or 3 hours per day for 5 days, 10-16 days (referred to as 14 days of self-administration), or 25-30 days (referred to as 30 days of self-administration) and could earn maximum of 100 infusions. A minimum of 10 remifentanil infusions for each self-administration day was set as a criterion whereas no criterion was used for saline self-administration.

### ‘Not Paired’ self-administration paradigm

In a subset of mice, acquisition involved responding on a fixed ratio 1 schedule for Ensure® alone for one day, then responding on a fixed ratio 1 schedule for remifentanil or saline alone for the remainder of self-administration (i.e. Not paired). These mice were also food deprived for initial lever training with Ensure®, then received food *ad libitum* throughout self-administration. Following initial lever training, self-administration was conducted as described above.

### *In vivo* activation of designer receptor exclusively activated by designer drugs

For DREADD studies, two to three days following completion of self-administration, a subset of mice received a bilateral intracranial infusion of AAV8-CamKII-hm3d(Gq)-mcherry (gift from Bryan Roth; Addgene viral prep #50476-AAV8; http://n2t.net/addgene:50476; RRID:Addgene_50476), AAV8-CamKII-hm4d(Gi)-mcherry (gift from Bryan Roth; Addgene viral prep #50477-AAV8; http://n2t.net/addgene:50477; RRID:Addgene_50477), or AAV8-Cam KII-GFP (UNC) into the prelimbic (coordinates from bregma: AP: +1.75, ML: ± 0.4; DV: −2.3). Mice were allowed to recover for five days, then were food deprived to 85-90% of their free feeding weight to be ran in the attention set-shifting task, as described below. Mice with viral infusions were given a saline intraperitoneal injection 30 minutes prior to testing in the visual cue test. On the day of the extradimensional shift, mice were given an intraperitoneal injection of clozapine-n-oxide (CNO; Tocris Biosciences, Cat. No: 4936). Mice with AAV8-Cam KII-hm3d(Gq)-mcherry were given 1.5mg/kg CNO, mice with AAV8-CamKII-hm4d(Gi)-mcherry were given 2.0mg/kg CNO, and mice with AAV8-CamKII-GFP were given either 1.5mg/kg or 2.0mg/kg to compare to both Gi or Gq mice and reduce the number of mice used. For *ex vivo* assessment of *in vivo* CNO, slices were prepared immediately following conclusion of behavioral testing or in a few instances, mice were given a second CNO injection on a subsequent day, and slices prepared at least 45 minutes following injection and recordings taken within 2 hours of slicing.

### Elevated plus maze

The morning following the last self-administration session, a subset of mice were tested for anxiety-like behaviors during a five minute test using the elevated plus maze (EPM; San Diego Instruments) as previously described^4^. Behavior was recorded using AnyMaze (Stoelting Co.) tracking software. Percent time in the open arms was calculated as total time in the open arms divided by total time in the maze.

### Forced swim test

A subset of mice were tested for immobility in a forced swim test approximately 16 days following the last day of self-administration, as previously described^4^. Briefly, mice were placed in an inescapable 800mL glass jar that was filled with 25°C water and were habituated for two minutes. Following habituation, time immobile was measured for four minutes using AnyMaze tracking software (Stoelting Co.). This test was performed following testing with the set-shifting task to avoid confounds of acute stress on subsequent behavior.

### Attention set-shifting task

Cognitive flexibility was measured using an operant based attentional set-shifting task that was modified based off of methods previously described^5^. Three to eight days following the last self-administration session, mice were food-deprived 85-90% of their free feeding weight. Training and testing were done in the operant conditioning chamber that was used for self-administration (Med Associates, Inc). During *food training*, a fixed ratio 1 schedule was used, whereby a right lever response (opposite lever to the reinforced lever during self-administration) resulted in delivery of a 50% liquid Ensure® reward. Food training sessions were 30 minutes in length and required mice to earn at least 50 rewards to progress to lever training. During *lever training*, levers were extended pseudorandomly with no more than two consecutive extensions of each lever. Mice were required to press the lever within 10 seconds of extension to receive a 50% liquid Ensure® reward, absence of which was deemed as an omission. Lever training consisted of a total of 90 trials (45 of each the left and right lever), with each followed by a 20 second time out. Mice had to reach criterion of five or fewer omissions on two consecutive days. Once lever training criterion was reached, *lever bias* was assessed through 7 trials, during which both the left and the right lever were reinforced.

The following day, *visual cue testing* was conducted until 150 trials or criterion was reached. Criterion was a streak of 10 consecutive correct responses. If this criterion was not reached on the first day of testing, a second or third day of testing was conducted. The correct response was the lever below the illuminated cue light and resulted in delivery of the Ensure® reward and a time out. Incorrect responses resulted in only the time out. Omissions were counted as stated above and were not counted towards a trial to criterion (i.e. neither correct nor error). The day following visual cue testing, *extradimensional shift testing* was conducted, and the reinforced lever was always the lever opposite of the lever bias; however, during this test the cue light was presented in a manner/order similar to that during the visual cue test. The day after criterion of the extradimensional shift was reached (i.e. 10 consecutive correct responses), reversal testing was conducted during which the reinforced lever was always that of the bias, however the cue light was presented identical to that used during the visual cue test. The criterion was still that mice had to reach 10 consecutive responses in a row. Once all attentional set-shifting tests were complete, food was provided *ad libitum* for the remainder of the experiments. For set-shifting experiments, data was excluded if >15 omissions in a given test was recorded as this may reflect mechanical issues or motoric issues. If these omissions were evident only during the reversal test, visual cue and extradimensional data was still used. Conversely, if these omissions were during the visual cue or extradimensional test, then all data was excluded as responding during these tests may have influenced responding during the extradimensional test or reversal test, respectively.

### Slice electrophysiology

Acute slice electrophysiology was performed either 24-72 hours, 14-21 days, or 40-45 days following the final self-administration session. In some cases, electrophysiology was conducted >21 days (for the 14-21 days group) if the mouse was tested in attention set-shifting and lever training took longer than average to acquire. Mice were mildly anesthetized with isoflurane (Henry Schein), decapitated, and the brain removed and put in ice-cold sucrose solution oxygenated using 95% O_2_ 5% CO_2_. A vibratome (Leica VT1000S) was used to obtain coronal slices (300μm) containing the mPFC. For CNO studies brains were removed at least 45 minutes following the systemic CNO injection and were sometimes conducted immediately following the extradimensional shift test and in other cases were conducted following a CNO injection in the home cage. Slices were immediately incubated at 31°C for 10 minutes in a solution 119mM NaCl, 2.5mM KCl, 1mM NaH_2_PO_4_, 26.2mM NaHCO_3_, 11mM glucose, 0.4mM ascorbic acid, 4mM MgCl_2_, and 1mM CaCl_2_. Slices were then removed, allowed to cool to room temperature and incubated further for a minimum of 35 minutes.

Whole-cell recordings were performed as previously described^4,6^. Briefly, slices were gravity perfused with oxygenated ACSF at a temperature of 29°C-33°C using at a flow rate of ~2-2.5 ml/minute. Sutter Integrated Patch Amplifier (IPA) with Igor Pro (Wave Metrics, Inc.) was used for the data acquisition software. Recordings were filtered at 2kHz and sampled at 5kHz for current-clamp recordings and voltage-clamp recordings assessing baclofen-evoked currents. Miniature postsynaptic current recordings were filtered at 2kHz and sampled at 20kHz.

Layer 5/6 pyramidal neurons were identified based on pyramidal-shaped soma, long apical dendrite extending superficially, capacitance (prelimbic >100pf; infralimbic >75pF), resting membrane potential (<−55mV), and lack of spontaneous activity^4,6-9^. For recordings in Gq+ pyramidal neurons, a resting membrane potential of <−50mV was used. For all recordings, adequate whole-cell access (Ra<40 MΩ) was maintained. Borosilicate glass pipettes were filled with 140mM K-Gluconate, 5.0mM HEPES, 1.1mM EGTA, 2.0mM MgCl_2_, 2.0mM Na_2_-ATP, 0.3mM Na-GTP, and 5.0mM phosphocreatine (pH 7.3, 290mOsm) for rheobase, action potential firing, and baclofen recordings. For both miniature and spontaneous excitatory and inhibitory postsynaptic currents, borosilicate glass pipettes were filled with 120mM CsMeSO4, 15mM CsCl, 10mM TEA-Cl, 8mM NaCl, 10mM HEPES, 5mM EGTA, 0.1mM spermine, 5mM QX-314, 4mM ATP-Mg, and 0.3mM GTP-Na. For miniature excitatory and inhibitory postsynaptic currents, 0.7mM lidocaine was added to the recording solution. mEPSCs were recorded at −72mV and mIPSC recorded at 0mV. To assess the influence of CNO bath application on rheobase, rheobase was measured as above followed by bath application of 5μM CNO and rheobase was tested 5-10 minutes following application.

### Histological assessment of virus expression

Accuracy and selectivity of virus targeting was assessed in slices prior to electrophysiology studies or post-hoc in fixed tissue. Viral placement was verified by fixing brains in 4% paraformaldehyde for 24-48 hours then placed in phosphate buffered saline until sliced using a vibratome during which 100μm were collected, mounted onto a slide using ProLong Gold Antifade Mountant (Thermo Fisher Scientific) and imaged using a Nikon inverted fluorescent microscope with a 4x objective. Virus placement was conducted for a subset of mice during electrophysiology recordings to assess the influence of CNO. Only mice with virus expressed bilaterally in the prelimbic were used in analyses. Animals with inaccurate targeting of virus or excessive virus expression outside the prelimbic were removed.

### Data analysis

2 (male, female) × 2 (saline, remifentanil) ANOVAs were used to test for statistical significance for initial rheobase studies, elevated plus maze, forced swim test, and attention set-shifting studies. Current-spike analysis was analyzed using a 2 (saline, remifentanil) way repeated-measures ANOVA. Responding across days in self-administration was analyzed using a repeated-measures ANOVA. Rheobase comparing only two groups (e.g. males versus females or paired versus not paired) was analyzed using an independent-samples t-test. For analysis of Gq-DREADD behavior and related recordings, a 2 (GFP, Gq) x 2 (saline, remifentanil) ANOVA was used. A t-test was used to compare attention set-shifting behavior and rheobase between saline mice with GFP and Gi-DREADD. For experiments where saline groups were combined a 2 (male, female) × 3 (saline, 14d remifentanil, 30d remifentanil) ANOVA was used. When interactions were found, main effects were not reported. Student-Newman-Keuls post-hoc comparisons were conducted when necessary. IPSCs/EPSCs were analyzed using MiniAnalysis software using a 5pA detection threshold (Synaptosoft). Data was analyzed using Sigma Plot (Systat Software, Inc) and graphed using Prism 7.0 (GraphPad). The threshold for statistical significance was p<0.05. Shapiro-Wilk test of normality and an equal variance assumption was used, when applicable. Statistical outliers were defined as being at least 2 standard deviations away from the mean and were removed from analyses. For electrophysiology studies, in most instances, no more than 3 recordings were used for a given metric (e.g., mEPSCs) from the same animal to reduce data bias. Exceptions were made when multiple recordings were acquired by different electrophysiologists on a given day.

### Data availability

The datasets generated during and/or analyzed during the current study are available from the corresponding author upon reasonable request.

